# TET2-driven activation of AGO2 links epigenetic remodeling to myeloid commitment and leukemia

**DOI:** 10.64898/2026.04.23.719229

**Authors:** Aleksey Lazarenkov, Gemma Valcárcel, Alba Martinez, Clara Berenguer, Anna V. López-Rubio, Mireia Obiols, Claudia Fontanet, Juan José Rodríguez-Sevilla, Gregoire Stik, Irmela Jeremias, Pablo Menendez, José Luis Sardina

## Abstract

DNA methylation dynamics shape hematopoietic differentiation and leukemogenesis; yet how the dioxygenase TET2—frequently mutated in myeloid malignancies—directs lineage-specific regulatory programs remains unclear. Here, we integrated DNA methylation, chromatin accessibility, 3D genome architecture, transcriptional profiling, and TET2 chromatin occupancy to define TET2-dependent control of human myeloid commitment. We found that TET2-bound regulatory regions gain short- and long-range chromatin interactions and that a distinct subset of distal, enhancer-enriched sites undergoes TET2-driven demethylation and activation. Among these, we identified an *AGO2* myeloid-specific intragenic enhancer that is frequently hypermethylated in TET2-mutant AML patients. *AGO2* expression stratifies patient survival, and AGO2 depletion abrogates leukemic engraftment *in vivo*. These findings uncover a TET2–AGO2 regulatory axis that integrates epigenetic remodeling, 3D genome reorganization, and leukemic fitness, and they highlight AGO2 as a potential biomarker and therapeutic target in myeloid leukemia.

## INTRODUCTION

DNA methylation (DNAm) at CpG dinucleotides is the main epigenetic modification in mammalian genomes and plays essential roles in development, differentiation and cancer ^1,2^. Although traditionally linked to transcriptional silencing, DNAm gain at certain promoters has been observed to coincide with increased gene expression, highlighting its context-dependent regulatory role ^3^. DNAm can be removed passively through the progressive dilution of methylation marks during DNA replication or actively via oxidation reactions catalyzed by the Ten-eleven-translocation (TET) family of dioxygenases^4^. In the active pathway, TET enzymes iteratively oxidize 5-methylcytosine (5mC) to 5-hydroxymethylcytosine (5hmC) and subsequent oxidized derivatives, which are then either lost through replication or enzymatically excised, ultimately restoring unmodified cytosine ^4^.

During blood cell fate decisions, the transcriptional impact of DNAm is shaped by the methylation sensitivity of lineage-instructive transcription factors (TFs). For example, Myc, Myb, and other bHLH TFs show reduced binding to methylated DNA, whereas KLF factors, such as KLF4, retain or even enhance binding in the presence of 5mC ^5^. Notably, KLF4, C/EBPα, and other pioneer myeloid TFs can recruit TET2 to cis-regulatory elements, linking methylation-insensitive TF binding to targeted demethylation during myeloid commitment ^6,7^. These observations thereby underscore the essential role of TET2-mediated enzymatic activity in modulating transcriptional outputs. *TET2* is among the most broadly expressed and recurrently mutated genes in myeloid malignancies, including acute myeloid leukemia (AML), myeloproliferative neoplasms (MPN), and myelodysplastic syndromes (MDS) ^8–10^, as well as in clonal hematopoiesis ^11,12^. Its loss-of-function drives differentiation arrest, myeloid bias, and aberrant inflammatory gene expression ^13–17^, yet the direct molecular mechanisms underlying these phenotypes remain poorly defined, mainly due to long-standing technical challenges in accurately mapping TET2 chromatin occupancy during hematopoietic differentiation.

To address this gap, we performed an integrative multi-omics analysis centered on TET2 binding during human myeloid commitment. Data revealed that TET2-bound regions undergo strengthening of both short- and long-range chromatin self-interactions during lineage specification, suggesting a role in higher-order genome organization. Of note, we also identified a discrete, enhancer-enriched subset of TET2 sites that become demethylated and chromatin-activated specifically during myeloid commitment, comprising 16 high-confidence targets. Among these, *AGO2* displayed the strongest methylation–expression response to TET2 depletion, driven by an intragenic enhancer that is selectively demethylated in the myeloid lineage and modulates *AGO2* expression, which in turn correlates with AML patient survival and leukemic potential *in vivo*.

## RESULTS

### A Flag-tag CRISPR knock-in strategy enables robust mapping of TET2 occupancy during myeloid commitment

Despite being a well-established facilitator of myeloid gene activation ^6,13,15,18,19^, the Cis Regulatory Elements (CREs) through which TET2 exerts this function remain poorly defined, largely due to technical challenges in mapping its precise genomic distribution. While TET2 occupancy has been characterized in terminally differentiated human myeloid cells ^20^, its function during myeloid lineage commitment remains largely unexplored. To address this critical gap, we employed a C/EBPα-driven myeloid commitment model that efficiently converts a B-cell leukemia line into functionally induced macrophages (iMac) ^21^. These iMacs recapitulate key features of primary macrophages, including robust phagocytic capacity, inflammasome competence, and responsiveness to inflammatory and pathogenic stimuli ^21–24^. To overcome the technical limitations to profile TET2, we introduced a Flag-tag at the *TET2* locus using CRISPR knock-in, generating tagged B-cells that were subsequently induced toward the myeloid fate **(Figure 1A)**. TET2 genomic occupancy was then profiled at the onset of the myeloid commitment (72 hours post-induction) in biological duplicates using a double-crosslinking protocol and anti-Flag immunoprecipitation followed by high-throughput sequencing (**Figure 1A**). Notably, at 72 hours post-induction, TET2 is significatively upregulated at both the mRNA and protein levels, coinciding with activation of the myeloid gene program (**Figures S1A-B**).

**Figure 1.**
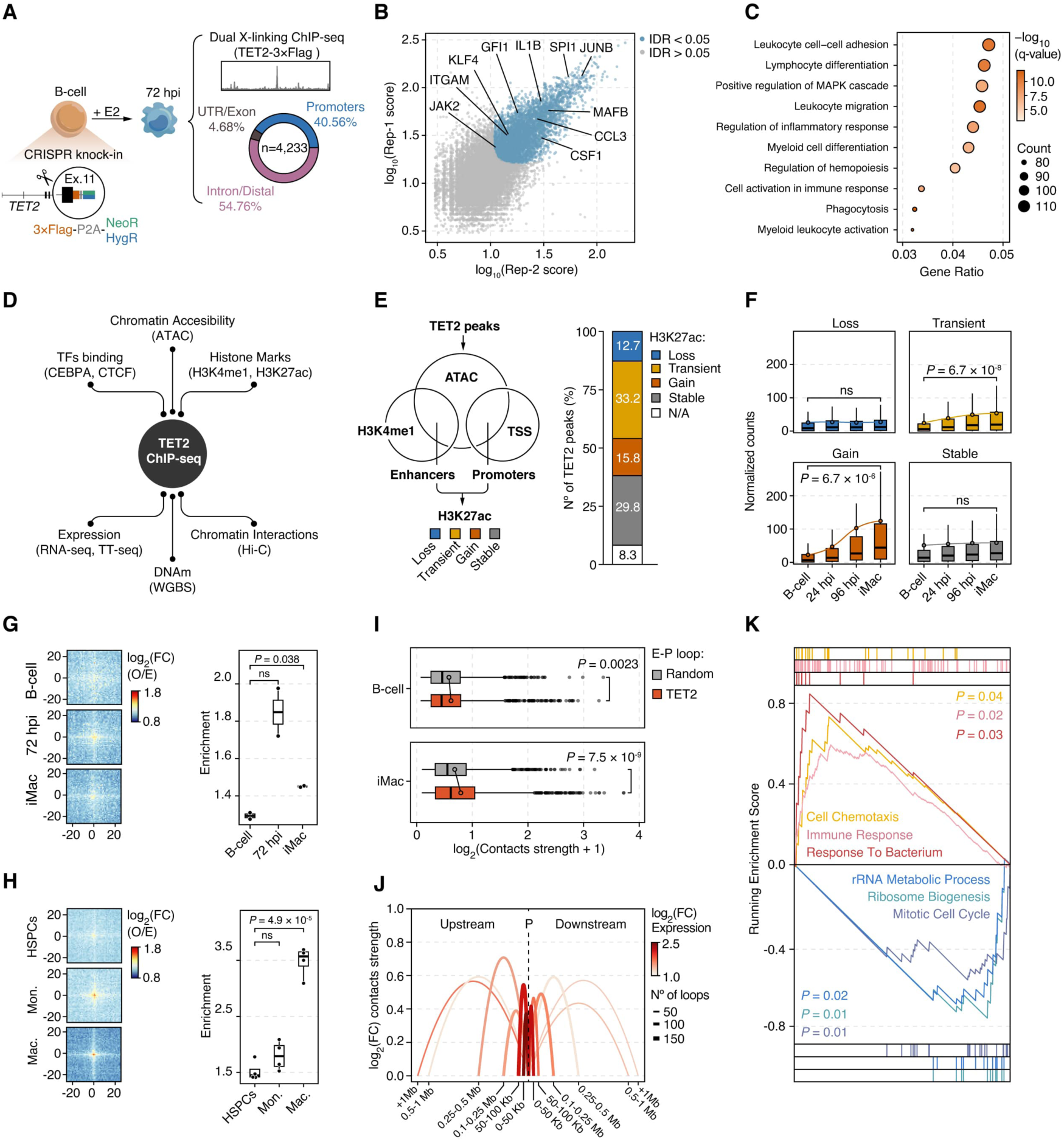
TET2 occupancy links enhancer–promoter architecture to activation of myeloid gene programs. (**A**) Schematic overview of the strategy used for TET2-3xFlag tagging and chromatin immunoprecipitation. Number of significant peaks and their annotation is shown. (**B**) Scatterplot of the IDR (Irreproducibility Discovery Rate) analysis showing common TET2 binding sites between the two biological replicates. (**C**) Gene Ontology (GO) enrichment analysis of genes associated with TET2 binding sites. Selected biological processes from the top 20 most significant GO terms are shown. (**D**) Schematic overview of TET2 genomic distribution data integrated with available datasets in the myeloid commitment cellular model ^23,24,26^. (**E**) Left: Schematic overview of the strategy used to classify TET2 binding sites into dynamic cis-regulatory elements (CREs). Right: Distribution of TET2-bound CREs according to their activation kinetics (H3K27ac levels). (**F**) Gene expression dynamics (by RNA-seq) during myeloid commitment for genes associated with the classified TET2-bound CREs. Dots with a stroke represent average values. (**G**) Left: Aggregate Peak Analysis (APA) plots showing long-range interactions (5-10 Mb) between TET2-bound regions during myeloid commitment. O/E: Observed over expected. Right: APA quantification showing the enrichment of the central bin versus the average number of contacts in all corners (iMac vs B-cell, n=2 per timepoint). (**H**) Left: APA plots showing long-range interactions (5-10 Mb) between TET2-bound regions during primary healthy myelopoiesis. O/E: Observed over expected. Right: APA quantification showing the enrichment of the central bin versus the average number of contacts in all corners (Mac. vs HSPCs, n=4 per timepoint). HSPCs: Hematopoietic Stem Progenitor cells; Mon.: Monocytes; Mac.; Macrophages. (**I**) Comparison of contact strength between random and TET2-bound enhancer–promoter (E–P) sites during myeloid commitment. (**J**) Arc diagram depicting average contact strength and differential expression of promoter-associated genes (iMac vs. B-cell) linked to TET2-bound E-P sites during myeloid commitment. (**K**) Gene Set Enrichment Analysis (GSEA) of genes associated with TET2-bound E-P sites (iMac vs B-cell). Selected significantly enriched terms are plotted.

Our chromatin occupancy analysis identified 4,233 highly confident TET2-binding sites (Irreproducibility Discovery Rate; IDR <0.05), including regions associated with key myeloid regulators such as *SPI1*, *KLF4*, *MAFB*, or *GFI1* **(Figures 1A-B).** In line with this, Gene Ontology analysis revealed that TET2-bound regions were enriched for genes related to myeloid commitment and macrophage functions **(Figure 1C).** Notably, most TET2 peaks localized to promoters and distal intra/intergenic elements (95.3%) (**Figure 1A, Figure S1C)** and were positioned either within 1 Kb of the TSS (29.2%) or at typical enhancer distances (10-100 Kb; 34.1%) from their nearest gene ^25^ (**Figure S1D)**, consistent with a role of TET2 occupancy in transcriptional regulation.

To refine the characterization of TET2 genomic distribution, we integrated multi-omics datasets capturing chromatin, transcriptional, and DNAm dynamics during myeloid commitment ^23,24,26^ **(Figure 1D).** We classified TET2-binding sites as promoters or enhancers based on chromatin accessibility and H3K4me1 enrichment, and assessed their activation by H3K27ac dynamics during myeloid commitment **(Figure 1E, left)**. Most TET2-bound regions (91.7%) overlapped with CREs, particularly those showing stable or transient H3K27ac gain during commitment **(Figure 1E, right; Figure S1E)**. Notably, these changes correlated with gene upregulation **(Figure 1F)**, highlighting TET2’s role in promoting myeloid gene expression.

Altogether, these results reveal extensive chromatin reconfiguration at TET2-bound regions during myeloid commitment, hinting at a coordinated reorganization of the underlying genomic landscape.

### TET2 binding sites undergo three-dimensional chromatin reorganization during myeloid commitment

To gain insight into chromatin dynamics of TET2-bound CREs during myeloid commitment, we integrated our occupancy data with genome-wide chromatin interaction maps (Hi-C) ^23^ **(Figure 1D)**. Motivated by recent reports suggesting TET2 forms biomolecular condensates ^27^, we investigated whether its binding is associated with changes in three-dimensional (3D) chromatin architecture. Remarkably, we observed a clear enrichment of long-range interactions (5–10 Mb) between TET2-bound sites during myeloid commitment **(Figure 1G)**. Notably, a similar pattern was detected during primary HSPC-to-macrophage differentiation **(Figure 1H)**, suggesting a potential role for TET2 in coordinating 3D chromatin rewiring. Importantly, C/EBPα, which highly co-localizes **(Figure S1F)** and interacts with TET2 ^6^, has also been reported to drive the formation of active 3D hubs through phase separation ^28^, suggesting a potential cooperative function of both proteins during myeloid commitment. Conversely, these TET2-associated 3D chromatin changes seem to occur independently of CTCF, as no changes in its occupancy were observed at TET2-bound sites during myeloid commitment **(Figure S1G)**.

While long-range contacts reflect higher-order chromatin organization, enhancer– promoter (E-P) loops directly link distal regulatory elements to gene transcription. To explore how TET2 contributes to local gene regulation, we analyzed TET2-bound E–P interactions. Using our previous E–P definition **(Figure 1E)**, we identified 1,358 loops occupied at both ends by TET2, which exhibited elevated contact frequency during myeloid commitment **(Figure S1H)**. Notably, TET2-bound enhancer–promoter (E–P) loops were significantly stronger than random pairs in iMacs (**Figure 1I**), indicating that TET2 binding may promote E–P association for transcriptional activation.

These loops were enriched in short-range contacts (<250 Kb) that correlated with gene upregulation during myeloid differentiation (**Figure 1J**). The associated transcriptional programs reflected activation of immune-related genes and repression of cell-cycle regulators, consistent with myeloid lineage acquisition **(Figure 1K).** To gain mechanistic insight, we next investigated whether TET2 catalytic activity contributes to loop establishment. We quantified DNAm levels from WGBS data **(Figure 1D)** at TET2 peaks within the E-P subset. Overall, DNAm changes did not significantly correlate with strengthened interactions during myeloid commitment (**Figure S1I**). Nonetheless, specific promoters and, to a greater extent, enhancers exhibited both demethylation and increased contact strength, as observed at key immune-related genes such as *RUNX3*, *IL3RA*, *IFNGR2,* or *NOD2* (**Figure S1J**).

Overall, while DNA demethylation seems insufficient to drive global E-P loop formation, locus-specific TET2 binding and associated demethylation may facilitate three-dimensional chromatin contacts that enable transcriptional activation of myeloid lineage-specific genes.

### An enhancer-enriched subset of TET2 binding sites is specifically demethylated and activated during myeloid commitment

We next examined DNAm dynamics across all TET2-bound regions to assess their impact on chromatin and transcriptional activity during myeloid commitment. Quantitative analysis revealed methylome remodeling from 24 hours post-induction (hpi) onward **(Figure 2A)**, coinciding with TET2 protein accumulation at 72 hpi **(Figure S1A)**. Notably, only ∼6% of the total DNAm signal was lost between B-cell and iMac stages **(Figure 2A)**, indicating that, despite widespread TET2 occupancy, demethylation is likely confined to a selective subset of CREs. To address this, we classified all TET2-binding sites as either Methylation Stable Peaks (MSPs) or Methylation Dynamic Peaks (MDPs), using a ΔDNAm > 0.15 threshold between the iMac and B-cell stages **(Figure 2B and Figure S2A)**. The distribution of the two groups was uneven: MSPs, showing an average DNAm loss of only 1.1%, accounted for 84.2% of all sites (3,427 regions), whereas MDPs, which lost >35% of DNAm, represented just 15.8% (642 regions) **(Figure 2B and Figure S2A)**. Consistently, MDPs exhibited significantly greater DNAm loss across all time points of myeloid commitment **(Figure S2B)**, suggesting them as a dynamically remodeled subset of TET2-bound regulatory elements.

**Figure 2.**
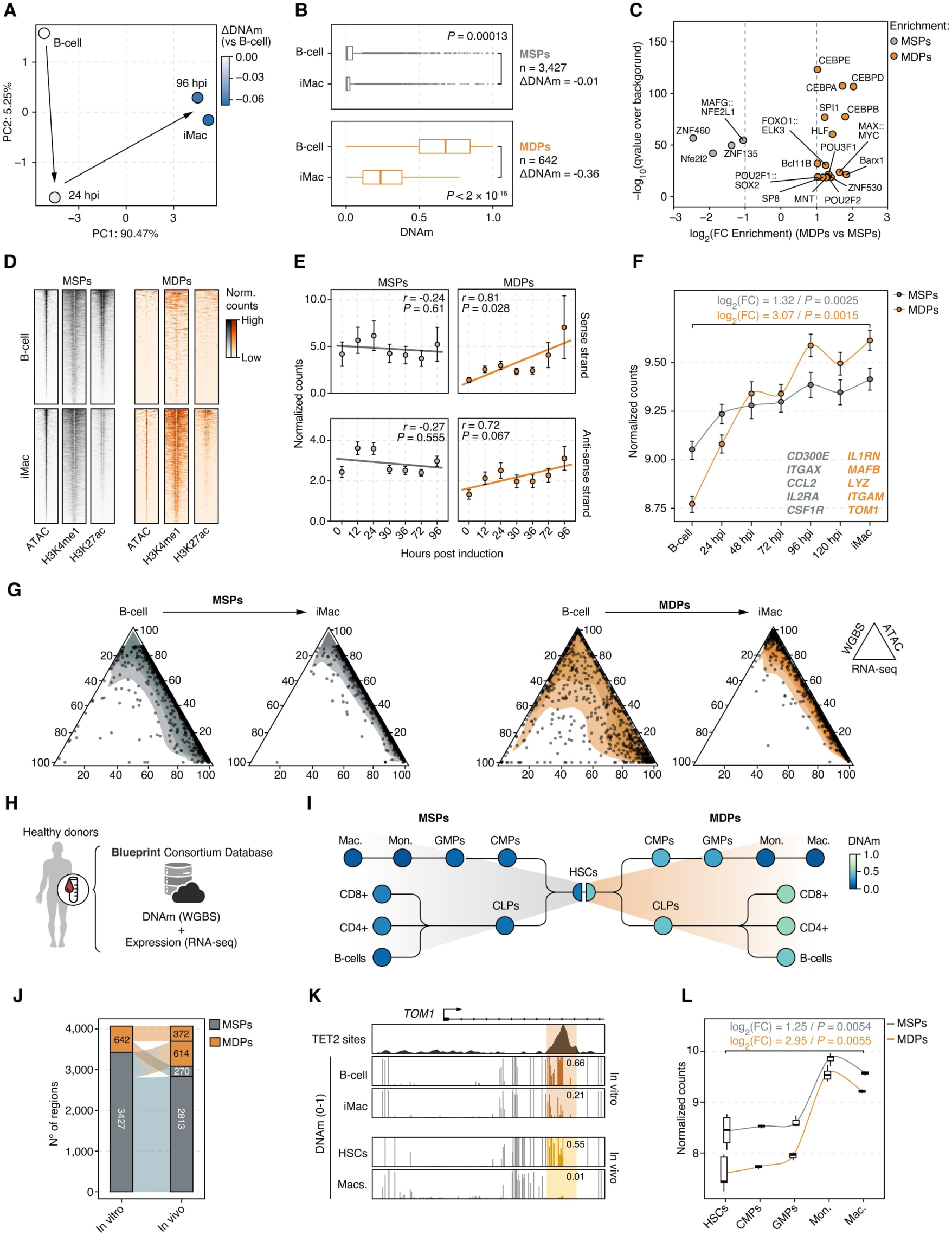
TET2 binding leads to DNA demethylation and activation of a highly dynamic subset of CREs during myeloid commitment. (**A**) Principal Component Analysis (PCA) of the DNAm dynamics at TET2 binding sites during myeloid commitment. Arrows indicate inferred DNAm trajectories. Color indicates the average DNAm changes during myeloid commitment (relative to B-cells). (**B**) Boxplots showing DNAm changes during myeloid commitment (iMac vs. B-cell) at TET2-bound Methylation Stable Peaks (MSPs) and TET2-bound Methylation Dynamic Peaks (MDPs). (**C**) Differential transcription factor (TF) enrichment analysis between MSPs and MDPs. (**D**) Genomic heatmaps showing chromatin accessibility (ATAC-seq) and activity (H3K4me1 and H3K27ac) during myeloid commitment at MSPs and MDPs. Signals are shown within a ±5 Kb window for each dataset. (**E**) Quantification of nascent RNA (TT-seq) dynamics at MSP and MDPs during myeloid commitment. Mean ± s.d. are shown for each timepoint (n = 2 per timepoint). The linear regression line is shown. P values were calculated from a two-sided Pearson correlation. (**F**) Gene expression dynamics (by RNA-seq) during myeloid commitment for genes associated with MSPs and MDPs. Mean ± s.d. are shown for each timepoint. Examples of genes from each group are shown in the bottom right. (**G**) Ternary plots showing changes in gene expression (RNA-seq), DNAm (WGBS) and chromatin accessibility (ATAC-seq) at MSPs and MDPs during myeloid commitment. Shaded regions depict point density. (**H**) Schematic overview of the human primary hematopoiesis samples and the expression (RNA-seq) and DNAm (WGBS) datasets analyzed in this study. Publicly available data from the Blueprint Consortium. (I) Bubble plot showing DNAm levels at MSPs and MDPs in primary samples during healthy hematopoiesis (n: HSCs = 9, CLPs = 3, CD8+ = 3, CD4+ = 3, B-cells = 3, CMPs = 3, GMPs = 3, Mon. = 9, Mac. = 6). Mon.; monocytes, Mac.: macrophages. (**J**) Comparison of the number of TET2-bound MSPs and MDPs defined in the “*in vitro*” myeloid commitment model (B-cell to iMac) versus those detected during primary HSC-to-macrophage differentiation “*in vivo*”. Shaded regions indicate reshuffling of the proportions between the groups. (**K**) Genome browser snapshot showing an intragenic MDP shared between the *in vitro* and *in vivo* myeloid commitment systems at the *TOM1* locus. Shaded region highlights the TET2 binding site. Average DNAm values within the region are shown. (**L**) Gene expression dynamics (by RNA-seq) during primary HSC-to-macrophage differentiation for genes associated with MSPs and MDPs. The solid line represents the average values. Mon.: monocytes; Mac.: macrophages.

To shed light on the distinct DNAm behaviors at TET2-bound regions, we next examined mechanistic differences between the two categories. TET2 binding strength was similar at MSPs and MDPs **(Figure S2C).** MSPs were enriched at CpG-dense promoter regions, whereas MDPs were biased toward distal sites (10–100 Kb from genes) and predominantly classified as enhancers **(Figures S2D-E)**. Next, we performed transcription factor (TF) motif enrichment analysis, revealing that MDPs were specifically enriched for key myeloid transcriptional regulators such as SPI1 and the CEBP family **(Figure 2C)**, including the known TET2 interactor CEBPA ^6^. Consistently, CEBPA occupancy was higher at MDPs during myeloid commitment **(Figure S2F)**, suggesting it may guide TET2 recruitment and demethylation at these sites. Regarding chromatin and transcriptional dynamics, only MDPs exhibited chromatin activation (measured by ATAC, H3K4me1, and H3K27ac) (**Figure 2D and Figure S2G**) and gained bi-directional nascent transcription (by TT-seq) during myeloid fate acquisition (**Figure 2E**), indicating full activation. In contrast, MSPs represent CREs already active in B cells (**Figures 2D-E and Figure S2G**). Accordingly, although genes associated with both TET2-bound CRE subsets were upregulated during myeloid commitment, activation was stronger for MDP-linked genes, which started from lower basal levels and reached higher expression than MSP-associated genes (log_2_(FC) = 3.07 vs. 1.32; respectively) (**Figure 2F**).

Overall, MSPs appear to correspond to promoter-proximal, poised elements that sustain basal myeloid identity, while MDPs are likely to characterize latent enhancers of inducible inflammatory programs (STAT1/ NF-κB-driven) **(Figure S2H)**. MDPs may thus represent genuine TET2-regulated CREs that undergo demethylation, increase accessibility, and promote strong transcriptional activation **(Figure 2G)**, offering a framework to evaluate their physiological importance in primary human myeloid differentiation.

### Enhancer-enriched TET2-bound regions show conserved demethylation and activation in primary human myeloid differentiation

To assess the physiological relevance of our findings, we analyzed DNAm dynamics at TET2-bound MSPs and MDPs across primary hematopoietic populations **(Figure 2H)**. MDPs showed pronounced, lineage-specific demethylation in myeloid cells (monocytes and macrophages) compared with HSCs, while remaining highly methylated in lymphoid populations. In contrast, MSPs were broadly unmethylated across all blood lineages **(Figure 2I)**. Notably, DNAm trajectories at MDPs converged between *in vitro* and *in vivo* myeloid commitment despite distinct starting states **(Figure S2I)**, indicating that our model faithfully recapitulates the myeloid-restricted demethylation of a TET2-regulated enhancer-enriched subset.

While these findings already underscore the physiological relevance of our TET2 bound MSP-MDPs dataset, we next aim to extend our characterization by comparing DNAm dynamics during *in vitro* and *in vivo* myeloid commitment. Applying the same ΔDNAm > 0.15 threshold used for the *in vitro* classification revealed some system-specific differences; however, more than half of the *in vitro* MDPs (372 out of 642) were conserved in the *in vivo* setting **(Figure 2J)**. These shared regions included CREs of myeloid-related genes such as the *TOM1* intragenic enhancer, which lost 45% and 54% of DNAm during *in vitro* and *in vivo* differentiation, respectively **(Figure 2K)**. Similar demethylation patterns were observed at a proximal *ITGAM* enhancer and at the *IL1RN* promoter **(Figure S2J)**. Consistently, transcriptional profiling of the *in vivo* MDP-associated genes mirrored the *in vitro* dynamics, with low basal expression in HSCs followed by marked upregulation in monocytes and macrophages compared to the MSP-linked genes **(Figure 2L)**. These changes paralleled the pronounced DNAm loss **(Figure 2I)** and the induction of TET2 expression during myeloid differentiation **(Figure S2K)**.

In summary, our comparative analysis of DNAm and transcriptional dynamics during *in vitro* and primary myeloid commitment reveals a conserved subset of TET2-bound, enhancer-enriched regions that undergo lineage-specific demethylation and robust gene activation, supporting their functional relevance and TET2’s potential role in orchestrating enhancer-driven myeloid gene programs.

### TET2 depletion disrupts lineage-specific DNA methylation dynamics during myeloid commitment

Having established that a subset of TET2-bound enhancers undergoes lineage-specific demethylation and activation during myeloid commitment, we next investigated to what extent these transcriptional changes depend on TET2’s catalytic activity. To directly assess TET2’s role in gene regulation during myeloid commitment, we generated three doxycycline-inducible *TET2* knockdown (TET2 iKD) models **(Figure 3A and Figure S3A)**. All iKDs exhibited a significant reduction in *TET2* expression (∼50%) following doxycycline (Dox) treatment, with shTET2.1 showing the strongest knockdown efficiency **(Figure S3A)**. Notably, shTET2.1 targets exon 3 of *TET2*—a region frequently mutated in myeloid malignancies ^13,29,30^ and was therefore selected for downstream characterization (hereafter referred to as shTET2). To ensure sustained TET2 silencing under Dox treatment, we validated shTET2 efficacy throughout myeloid commitment **(Figure S3B)**. Consistently, shTET2 cells showed markedly lower 5hmC levels **(Figure S3C)**, confirming reduced TET2 catalytic activity. With the shTET2 model validated, we next examined how loss of TET2 impacts gene expression and DNAm at TET2-bound regulatory elements during myeloid commitment **(Figure 3A)**. DNAm analyses revealed a progressive gain during myeloid commitment **(Figure 3B and Figure S3D)**, culminating at the iMac stage with 1,794 differentially hypermethylated positions and an average DNAm increase of 21.3% **(Figure S3D)**.

**Figure 3.**
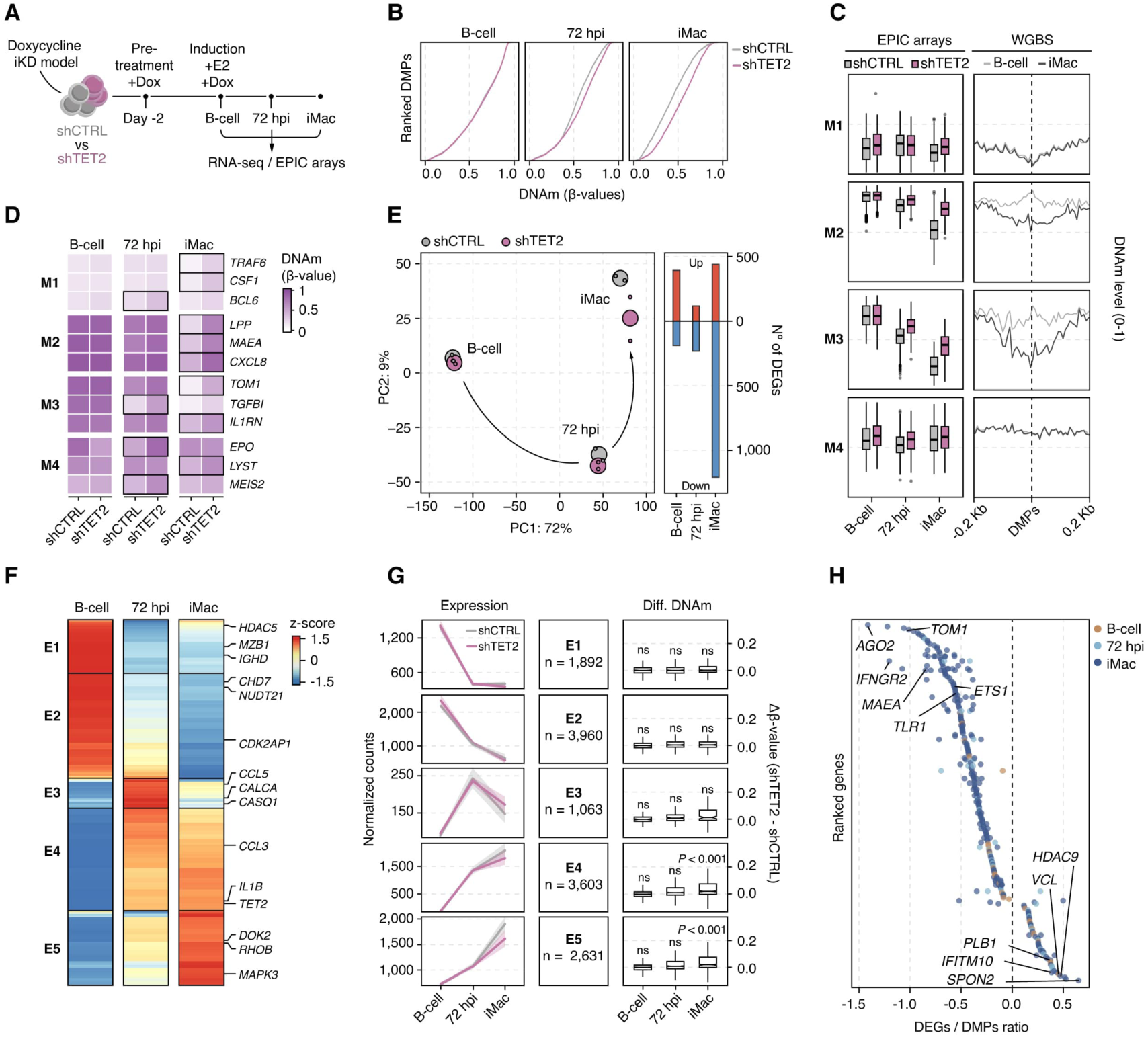
TET2 depletion impairs DNAm-mediated activation of myeloid gene programs. (**A**) Schematic overview of the integrative experimental design used to assess the impact of TET2 depletion on DNAm and transcriptional dynamics during myeloid commitment. (**B**) DNAm (β-values) of ranked Differentially Methylated Positions (DMPs) detected in shTET2 vs. shCTRL cells during myeloid commitment. (**C**) Left: Quantification of DNAm (β-values) for DMPs within DNAm clusters (M1-4) during myeloid commitment. Right: Average DNAm (WGBS) profiles levels in B cells and iMacs centered on DMPs from each cluster (M1-4). (**D**) Heatmap of DNAm levels (β-values) during myeloid commitment for DMPs hypermethylated in TET2-depleted cells, grouped by DNAm cluster, with associated gene names shown. Significant changes (p < 0.01) are indicated by a black outline. (**E**) Left: Principal Component Analysis (PCA) of transcriptome dynamics during myeloid commitment in shTET2 and control cells. Small dots represent individual biological replicates (n = 2), and large dots indicate the average of each timepoint. Arrows indicate inferred trajectories of myeloid commitment. Right: Number of significant up- or down-regulated genes in shTET2 cells at each timepoint. (**F**) Heatmap showing the clustering of transcriptional dynamics (E1-5) for the most significant gene expression changes during myeloid commitment in control conditions (FDR < 0.05, absolute log_2_(FC) > 1). (**G**) Evaluation of DNAm-expression dynamics in expression clusters (E1-5) during shTET2 versus control myeloid commitment. Left: Quantification of gene expression in each cluster; mean ± s.d. are shown for each timepoint. Solid lines represent the average value and shaded regions indicate the standard deviation (n=2 per timepoint). Right: DNAm changes associated with genes in each cluster. Only probes within DNAm-dynamic regions (n = 11,860), as defined by Valcarcel et al. ^26^, were included in the quantification. (**H**) Ranked ratio of overlapping DEGs and DMPs identified in shTET2 versus shCTRL cells during myeloid commitment. Selected genes with significant hypermethylation and downregulation are highlighted on the left side of the dashed line, while those with hypermethylation and upregulation are shown on the right.

To gain a deeper understanding of how TET2 depletion influences DNAm dynamics during myeloid commitment, we clustered all differentially methylated positions (DMPs) into four groups exhibiting distinct temporal trends **(Figure S3E)**. Only DMPs within the M2 and M3 clusters displayed DNAm loss during the process **(Figure 3C, left)**. These loci overlapped with broader regions (±200 bp) undergoing demethylation during myeloid commitment **(Figure 3C, right)**, suggesting that the M2 and M3 clusters reflect coordinated regulatory domains rather than isolated DNAm events. Consistent with TET2’s catalytic role, shTET2 cells showed pronounced hypermethylation within M2–M3 clusters at 72 hours post-induction (hpi) and at the iMac stage **(Figure 3C, left)**. Genes associated with these clusters were enriched for immune response and myeloid activation pathways **(Figure S3F)**. Despite these differences, all clusters contained key regulators of macrophage differentiation and function—including *CXCL8*, *IL1RN*, *CSF1*, and *EPO* in M1 through M4, respectively **(Figure 3D)**.

Altogether, these findings reveal that TET2 loss disrupts the coordinated demethylation of lineage-associated CREs during myeloid differentiation, particularly at loci involved in immune activation and myeloid cell function, thereby establishing a methylation landscape poised to influence transcriptional output.

### TET2 depletion impairs activation of myeloid-specific gene expression programs through defective DNA demethylation

Building on the observed DNAm changes, we next interrogated transcriptional alterations in TET2 iKD cells by RNA-seq. Mirroring the progressive changes in DNAm, the majority of differentially expressed genes (DEGs) were detected at the iMac stage (**Figure 3E and Figure S3G**). Notably, principal component analysis (PCA) indicated an overall delay in the transcriptional dynamics of myeloid differentiation in shTET2 conditions **(Figure 3E, left)**, accompanied by widespread gene silencing (n = 1,208 DEGs) at the iMac stage **(Figure 3E, right; Figure S3G)**. Among the downregulated genes were key myeloid regulators such as *MAF*, *IL1B*, and *IRF8* **(Figure S3G)**, collectively associated with phagocytosis, immune response, and differentiation processes **(Figure S3H)**.

To gain deeper insight into TET2’s role in shaping normal transcriptional dynamics during myeloid commitment, we applied a clustering approach to define transcriptionally dynamic gene groups during the process and assessed how these patterns were altered upon TET2 depletion. Five transcriptional clusters (E1–E5) were identified **(Figure 3F)**, corresponding to the silencing of B-cell–related programs (E1) and DNA replication and chromatin organization pathways (E2), transient activation of calcium and phospholipase signaling genes (E3), and induction of immune and myeloid cell activation, leukocyte migration, and vesicle transport processes (E4–E5) (**Figure 3F and Figure S3I)**. Integration of transcriptional and DNAm profiles revealed that, while clusters E1–E2 showed minimal changes, E3–E5 were markedly affected by TET2 loss. At the transcriptional level, shTET2 cells displayed impaired upregulation of E4–E5 genes and, unexpectedly, higher expression of E3 genes compared to controls **(Figure 3G, left)**. Consistent with these observations, hypermethylation tendencies were detected within the affected clusters, reaching significance at the iMac stage in E4 and E5 **(Figure 3G, right)**. This relationship was further supported by analyzing the overlap between TET2-dependent DMPs and DEGs, where promoter or genic hypermethylation was predominantly associated with gene silencing in TET2-depleted conditions **(Figure 3H)**. Notably, this included genes such as *TOM1*, *IFNGR2*, and *IL1RN* **(Extended Data Figure 3J)**, which we previously identified as TET2-bound methylation dynamic CREs during myeloid commitment **(Figure 2K and Figure S2J)**.

Altogether, these findings highlight the importance of combining TET2 chromatin occupancy data with DNAm and transcriptional analyses to precisely identify its direct targets and to understand how their regulation influences myeloid identity.

### Integrative multi-omics analyses reveal AGO2 as a direct TET2 target in myeloid commitment

To identify direct TET2 targets, we intersected TET2-bound MDPs–defined as TET2 peaks (ChIP–seq) overlapping WGBS-demethylated regions (≥15% DNAm loss between B-cell and iMac stages)– with TET2 iKD DEGs, DMPs **(Figure 4A)**. This analysis yielded 16 candidate genes, including the interferon-gamma receptor *IFNGR2*, the histone deacetylase HDAC9, and other genes of interest, such a *TOM1*, *NDRG1*, and *AGO2*. Notably, AGO2, a key RNAi regulator, showed the strongest DNAm–expression dependency in TET2-depleted cells **(Figure 3H and Figure S3J)**, uncovering a previously unrecognized role for TET2 in modulating crucial components of the RNA interference pathway.

**Figure 4.**
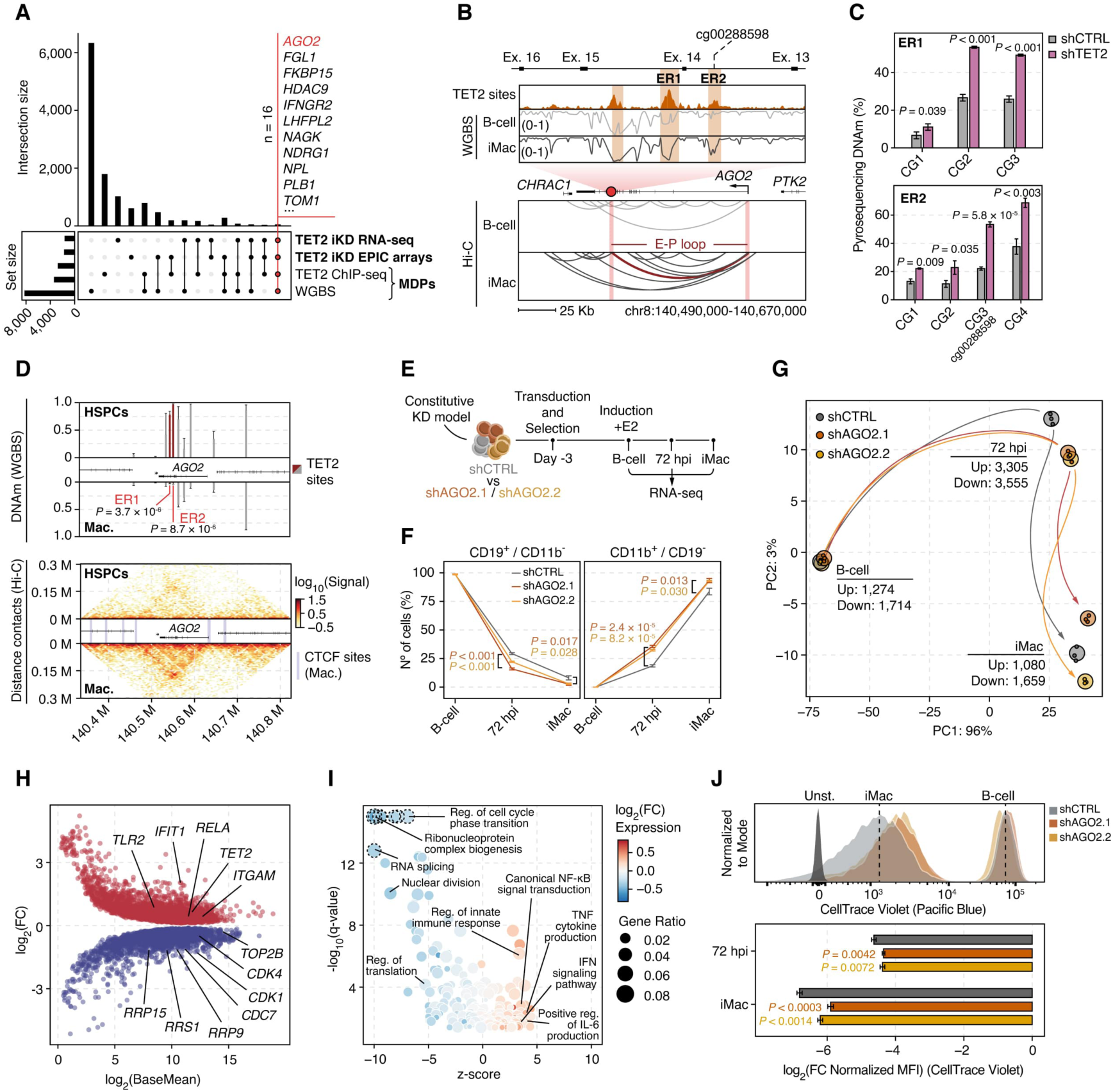
Integrative multi-omics analysis identifies AGO2 as a TET2-dependent regulator of myeloid commitment. (**A**) UpSet plot showing the overlap between TET2-related datasets during myeloid commitment, including chromatin occupancy, DNA-demethylated regions (WGBS) ^24^, and DMPs / DEGs in TET2-depleted cells. Candidate genes from the full overlap are indicated in **Table S1**. MDPs: Methylation Dynamic Peaks (as defined in Figure 2). (**B**) Genome browser snapshot of the *AGO2*-associated region of interest. Top: Close-up genomic tracks showing TET2 binding (shadowed) and DNAm dynamics during myeloid commitment. TET2-bound enhancer regions with DNAm loss (ER1 and ER2) and the DMP of interest are annotated. Bottom: Hi-C interaction arc diagram at the *AGO2* locus in B-cell and iMac cells. Only interactions with a score > 1 and a distance > 20 Kb are shown. The red arc depicts the iMac-specific loop connecting the region of interest and the *AGO2* promoter. (**C**) Quantification of DNAm by pyrosequencing at ER1 and ER2 regions in TET2-depleted and control iMacs. Mean ± s.d. are shown for each CpG (n = 3). (**D**) Genome browser snapshot of the *AGO2* locus in primary human HSPCs and macrophages. Top: DNAm levels at TET2 binding sites within the *AGO2* locus. Mean ± s.d. are shown for each region (n: HSPCs = 4, Mac. = 4). Bottom: Hi-C contact heatmap depicting changes in contact strength between HSPCs and macrophages. Mac.: macrophages. (**E**) Schematic overview of the experimental design used to assess the role of *AGO2* in myeloid commitment. (**F**) Flow cytometry analysis of cell surface markers in shAGO2-depleted and control cells. Percentages of B-cells (CD19^+^/CD11b^-^) and iMac (CD11b^+^/CD19^-^) are shown. Mean ± s.d. are shown for each timepoint (n = 3). (**G**) Principal Component Analysis (PCA) of transcriptome dynamics during myeloid commitment in shAGO2 and control cells. Small dots represent individual biological replicates (n = 3), and large dots indicate the average for each timepoint. Arrows indicate inferred trajectories of myeloid commitment. (**H**) MA plot showing DEGs in shAGO2 cells at 72 hpi. (**I**) Gene Ontology (GO) enrichment analysis of DEGs in shAGO2 cells at 72 hpi. Over- and under-represented biological processes (z-score) are plotted, with color indicating the average log_2_(FC) of genes within each term. Dotted outlines indicate outlier terms (-log_10_(q-value) > 15). (**J**) Proliferation assay by flow cytometry in shAGO2 cells during myeloid commitment. Top: representative histograms showing CellTrace Violet signal. Bottom: quantification of CellTrace signal normalized to the B-cell. Mean ± s.d. are shown for each timepoint (n = 3). MFI: Mean Fluorescence Intensity. Unst: unstained control.

To further explore this regulatory mechanism, we next examined the *AGO2* locus and observed extensive TET2 binding across its gene body, including the intragenic region identified in the multi-omics overlap (DMP: cg00288598) **(Figure 4B, top; Figure S4A)**. This region displayed features of enhancer activation, marked by gain of H3K4me1 and H3K27ac during myeloid commitment and CEBPA recruitment from 24 hpi onward **(Figure S4A)**. Chromatin conformation analysis further revealed increased contacts at the *AGO2* locus, including enhancer–promoter looping between the candidate region and the *AGO2* promoter **(Figure 4B, bottom)**, consistent with chromatin remodeling and transcriptional changes. Notably, H3K27ac enrichment peaked transiently at 72 hpi, coinciding with expression data **(Figures S4A, S3J)**, suggesting that *AGO2* is transiently upregulated at intermediate differentiation stages via TET2-dependent DNA demethylation before declining in more committed cells. Importantly, the region of interest encompassed multiple TET2 binding sites that underwent demethylation during myeloid commitment, hereafter referred to as enhancer regions ER1 and ER2. The cg00288598 DMP, located within ER2, displayed clear hypermethylation in TET2 iKD cells. To validate these findings, we quantified DNAm levels at three additional CpGs per site by targeted pyrosequencing, detecting significant ER1/ER2 hypermethylation in iMacs with reduced TET2 levels **(Figure 4C)**, confirming strong catalytic activity at this locus. We next examined publicly available WGBS and Hi-C datasets from healthy myelopoiesis to assess the physiological relevance of our findings. Consistent with our model, ER1 and ER2 were the only TET2-bound sites exhibiting significant demethylation in macrophages compared with hematopoietic progenitors **(Figure 4D, top)**. Likewise, chromatin interaction data revealed the macrophage-specific formation of a chromatin domain at the AGO2 locus, anchored by CTCF sites **(Figure 4D, bottom)**.

In summary, our integrative analysis of TET2 binding, DNAm and transcriptional data revealed novel catalytic targets potentially involved in myeloid commitment. Among these, an *AGO2* enhancer exhibited TET2-dependent demethylation, accompanied by coordinated transcriptional and chromatin changes during human *in vitro* and primary myelopoiesis.

### AGO2 downregulation promotes myeloid commitment while restraining cell proliferation

Building on our identification of *AGO2* as a direct TET2 target during myeloid commitment and on previous studies suggesting a role for *AGO2* in hematopoiesis ^31,32^, we sought to investigate its functional role in this process. To this end, we generated two constitutive *AGO2* knockdown cell lines (shAGO2.1 and shAGO2.2), validated efficient depletion, and characterized their behavior upon induction of myeloid commitment **(Figures 4E and Figure S4C)**. Remarkably, *AGO2* knockdown accelerated myeloid cell fate acquisition, as evidenced by faster loss of the B-cell marker CD19 and gain of the myeloid marker CD11b (flow cytometry) **(Figure 4F)**, and by principal component analysis of the transcriptional landscape (RNA-seq) **(Figure 4G)**. Despite minor differences in depletion efficiency **(Figure S4C)**, both shAGO2.1 and shAGO2.2 lines shared over 60% of differentially expressed genes **(Figure S4D)**, showing common downregulation of metabolism, translation, and cell cycle programs, and upregulation of inflammation-related pathways, including interferon signaling **(Figure S4E)**. Specifically, ribosome and rRNA metabolism genes, such as *RRP15*, *RRP9*, and *RRS1*, were consistently downregulated across all timepoints, while mitotic genes (*CDK1*, *CDK4*, *CDC7*) were affected primarily at the iMac and 72 hpi stages. The upregulated gene set included a broad range of immune response genes, such as *TLR2*, *IFIT1*, *ITGAM*, and *IL1B* **(Figures 4H–I and Figure S4G)**, the latter being elevated at all timepoints, highlighting the potential impact of reduced *AGO2* levels on myeloid rewiring.

We next investigated whether the observed *AGO2* knockdown–induced transcriptional silencing of mitotic regulators translated into phenotypic changes. Proliferation analysis using CellTrace Violet dye dilution revealed that AGO2-depleted cells slowed their cell cycle more rapidly than control cells, evident at 72 hpi and in iMacs **(Figure 4J)**. Notably, a similar effect was observed even at the uninduced B-cell stage, with proliferation markedly reduced as early as three days post-seeding **(Figures S4H–I)**.

Together, these results suggest that AGO2 reduction slows highly proliferative processes, thereby facilitating the transition toward non-dividing, fully committed myeloid cells.

### AGO2 locus methylation status predicts expression and prognosis in AML

The observed effects of AGO2 on myeloid commitment and proliferation suggested that it may also contribute to leukemogenesis, consistent with previous reports hinting at a similar role ^33^. To explore this, we analyzed DNAm and expression data from publicly available adult AML patient datasets **(Figure 5A)**.

**Figure 5.**
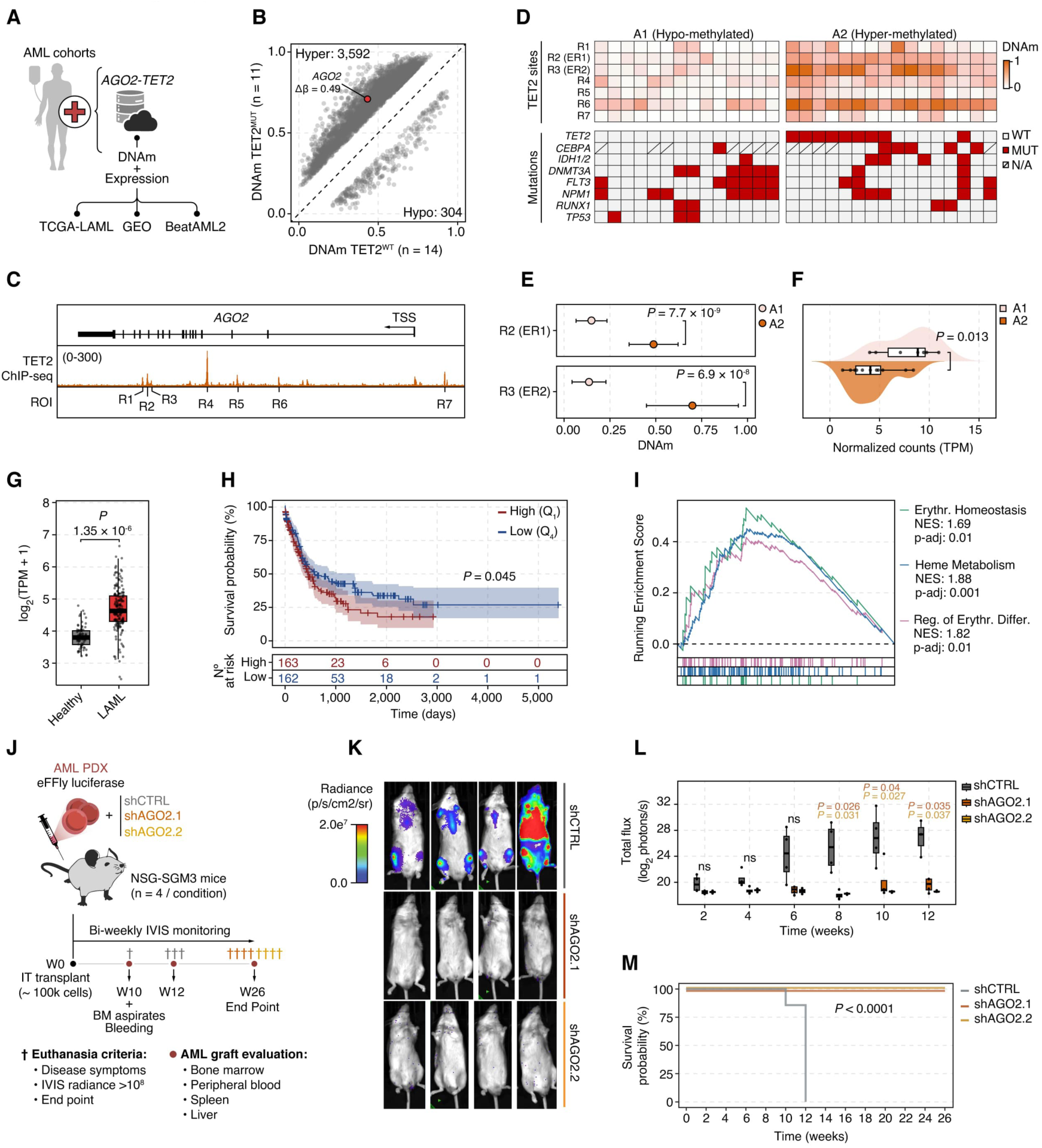
AGO2 is hypermethylated in TET2^MUT^ AML patients and its silencing impairs *in vivo* leukemic progression. (**A**) Schematic overview of the human primary AML samples and the expression (RNA-seq) and DNAm (WGBS) datasets analyzed in this study. Publicly available data from TCGA-LAML, GEO, and BeatAML2. (**B**) Scatter plot showing DMPs between TET2^MUT^ and TET2^WT^ AML patients. (**C**) Genome browser snapshot illustrating the AGO2 Regions of Interest (ROI; R1-R7) defined by TET2 chromatin occupancy. (**D**) Evaluation of AGO2 DNAm status in AML patients (n = 30). Top: Heatmap showing patient clustering (A1-A2) based on DNAm levels at *AGO2* ROI (R1-R7). Bottom: Heatmap showing the mutational profile of each patient. N/A: indicates not profiled for the specific mutation. (**E**) Quantification of DNAm levels at ER1-ER2 sites across AML clusters (A1-2). Mean ± s.d. are shown for each condition. (**F**) Violin plot showing AGO2 expression in AML cluster A1 (n = 7) and A2 (n = 11) from available patient samples. (**G**) Boxplots showing mRNA *AGO2* levels in patient samples from the TCGA-LAML cohort (n = 173) compared with healthy bone marrow samples from the GTEx dataset (n = 70). Data from GEPIA2 portal ^49^. (**H**) Kaplan-Meier survival plots for patients with high (Q_1_: Quartile 1; n = 163) versus low (Q_4_: Quartile 4; n = 162) *AGO2* expression in the BEAT-AML2 cohort. Shaded areas indicate the 95% confidence interval. Statistical significance was assessed using the Mantel-Cox Log-rank test. Risk table is shown below. (**I**) Gene Set Enrichment Analysis (GSEA) of the DEGs in high (Q_1_) vs. low (Q_4_) *AGO2* patients from the BEAT-AML2 cohort. Selected erythroid-related terms with significant enrichment are plotted. NES: Normalized Enrichment Score. (**J**) Schematic overview of the experimental design used to evaluate AGO2 depletion effect on leukemia onset and development using an AML-PDX mouse model. (**K**) Whole-body in vivo imaging (IVIS) showing radiance signal at 10 weeks post-injection in mice injected with shAGO2 or shCTRL AML-PDX cells. (**L**) Quantification of IVIS radiance from week 2 post-injection to week 12, when all mice injected with shAGO2 AML-PDX cells reached the experimental endpoint. (**M**) Kaplan-Meier survival plots of mice injected with shAGO2 or shCTRL AML-PDX cells. Statistical significance was assessed using the Mantel-Cox Log-rank test.

First, to investigate the impact of TET2 mutations on AGO2 DNAm in AML, we analyzed EPIC array data from the TCGA-LAML cohort, comparing TET2^MUT^ and TET2^WT^ samples **(Figure 5B)**. Consistent with our observations during myeloid commitment, TET2^MUT^ samples exhibited widespread hypermethylation, notably including the same *AGO2* cg00288598 DMP that became hypermethylated upon *TET2* knockdown in the vitro model **(Figure 5B)**. To extend these findings, we examined DNAm across all significant TET2 binding sites within the *AGO2* locus (regions of interest -ROIs-: R1–R7) **(Figure 5C)** using WGBS data from AML patients harboring common mutations. Unsupervised clustering revealed two groups with hypo-(A1) and hypermethylated (A2) profiles **(Figure 5D)**. Notably, all TET2^MUT^ samples (n = 8/8) fell into the hypermethylated A2 group, confirming the association between TET2 loss and increased DNAm at this locus. In contrast, the A1 group was enriched for DNMT3A^MUT^ samples (n = 6/8), highlighting the locus’s DNAm dependency **(Figure 5D)**. Importantly, quantification of *AGO2* ROIs during healthy hematopoiesis confirmed that R2–R3, corresponding to ER1–ER2 **(Figure 4B)**, were specifically demethylated during monocyte differentiation **(Figures S5A-B)**. Notably, these two sites were the most differentially methylated between the A1 and A2 AML clusters **(Figure 5E),** and hypermethylation of the *AGO2* cg00288598 probe (within ER2) correlated with poorer overall survival **(Figure S5C)**, likely reflecting the aggressive nature of immature blasts.

Finally, DNAm status correlated with gene expression, as the hypermethylated A2 cluster exhibited lower levels of *AGO2* expression **(Figure 5F)**, supporting the proposed methylation–expression association and providing a rationale to investigate AGO2’s role in AML.

### AGO2 levels regulate AML transcriptional programs and in vivo leukemogenesis

Having established a link between *AGO2* methylation and expression in AML, we next examined how *AGO2* expression levels relate to leukemic phenotypes and patient outcomes. First, we observed that, compared to healthy counterparts, AML samples exhibited elevated *AGO2* expression **(Figure 5G)**, suggesting that gene upregulation may contribute to leukemic maintenance. Indeed, overall survival analyses revealed a significantly worse prognosis in patients with high *AGO2* expression **(Figure 5H)**. Next, RNA-seq analysis of AML patients with high versus low *AGO2* expression revealed that *AGO2* upregulation was associated with repression of proliferation regulators (*TP53*, *TP63*) and activation of erythroid differentiation factors (*TAL1*, *EPOR*) **(Figure 5I and Figure S5D)**. This transcriptional signature aligns with *AGO2*’s expression specificity **(Figure S5E)** and its reported role in murine erythropoiesis ^32,34^; and may contribute to leukemic bias, as high-AGO2 samples were enriched in M6 (acute erythroid) and M7 (acute megakaryoblastic) AML subtypes **(Figure S5F)**. Conversely, NPM1-mutated patients were enriched among the low-AGO2 group, which consistently displayed lower *AGO2* expression compared to NPM1^WT^ AML cases **(Figure S5G–H)**. This relationship might influence disease progression, as AGO2 and NPM1 have been reported to physically interact with each other and with microRNAs ^35–37^. Altogether, these associations suggest that *AGO2* levels may influence leukemic lineage bias and disease progression in different leukemic settings.

Finally, to functionally assess whether *AGO2* depletion affects leukemogenic capacity *in vivo*, we used a patient-derived xenograft (PDX) leukemia model. Aggressive AML blasts (AML-579) ^38,39^ were transduced with lentiviruses expressing either control shRNA (shCTRL) or shRNAs targeting *AGO2* (shAGO2.1 and shAGO2.2). *AGO2* knockdown resulted in marked protein reduction **(Figure S5I)**, and the cells were subsequently injected into recipient mice for *in vivo* imaging (IVIS) and endpoint tissue analysis **(Figure 5J)**. Remarkably, AGO2 depletion completely abrogated tumor development, whereas control mice exhibited elevated IVIS radiance and leukemic burden at 10 weeks post-injection (wpi) **(Figure 5K and Figure S5J)**. Bone marrow (BM) aspirates revealed a higher percentage of human AML blasts (HLA⁺) in controls, which was also detected in peripheral blood (PB), to a lesser extent **(Figure S5K)**. Bi-weekly IVIS monitoring revealed robust engraftment in control mice from 6 wpi until the 12-week endpoint, whereas AGO2 depleted mice remained healthy for over six months **(Figures 5L-M)**. Consistently, at euthanasia, only control animals showed detectable levels of human blasts in BM, liver, and spleen **(Figure S5L)**, along with classic AML features such as splenomegaly **(Figure S5M)**.

Taken together, these results highlight a key role for AGO2 in AML, linking its expression to transcriptional programs in patients and demonstrating that AGO2 depletion effectively prevents leukemic engraftment *in vivo*.

## DISCUSSION

In this study, we applied an integrative multi-omics framework combining DNA methylation, chromatin activity and conformation, transcriptional data, a newly generated TET2 binding map, and functional studies to delineate TET2-mediated regulatory programs during myeloid commitment. This approach revealed an enhancer-enriched subset of high-confidence TET2 targets, including a key regulatory element at the *AGO2* locus that undergoes lineage-specific demethylation and activation—epigenetic processes that become disrupted in AML patients. Notably, AGO2 depletion impaired leukemia development *in vivo*, highlighting a functional axis with potential prognostic and therapeutic relevance.

By Flag-tagged pulldown in a physiological model, we reveal TET2 chromatin occupancy during human myeloid commitment, capturing a transitional stage not previously characterized in studies of fully differentiated myeloid cells ^20^. This approach showed that TET2 is recruited early during myeloid commitment, likely preceding widespread DNA methylation remodeling, highlighting its dynamic role in fate specification. Integration with complementary epigenomic and transcriptomic datasets revealed a pronounced TET2 bias toward regulatory elements—not only enhancers, as previously described ^40^, but also promoters of key myeloid genes, including *KLF4*, *ITGAM*, and *IL1B*—where binding correlated with stable or transient H3K27ac deposition and gene activation. Notably, we showed that TET2-bound promoters are largely unmethylated across the human hematopoietic system, making 5hmC an unreliable proxy for TET2 binding and underscoring the need for direct mapping to define TET2’s role in cell fate specification.

The presence of TET2 at unmethylated promoters suggests roles beyond demethylation during myeloid commitment. Notably, TET2 may shape chromatin architecture independently of DNAm, strengthening both local loops and long-range interactions (5–10 Mb) at its binding sites. This effect could involve biomolecular condensation of TET2 ^27^ and cooperative interactions with factors such as CEBPA ^28^. Previous studies in the murine hematopoietic system have highlighted distinct biases induced by the loss of TET2 catalytic activity versus complete protein loss ^15^, supporting the idea that non-catalytic functions may have physiological relevance. However, the precise mechanisms of this non-catalytic remodeling and its functional implications in human myeloid commitment remain to be defined.

Conversely, we identified a discrete set of enhancer-enriched TET2 sites (MDPs, Methylation Dynamic Peaks) that acquired chromatin accessibility and active histone concurrent with DNA demethylation during myeloid commitment, resulting in robust upregulation of their associated genes. These genes are enriched for inflammatory and stimulus-responsive functions and may be regulated by the STAT1 and RELA/NFKB1 pathways. While the temporal order of demethylation and activation remains unresolved, our previous work demonstrated that CRISPR-dCas9-mediated DNA demethylation can trigger gene activation, as exemplified by *IL1RN* ^24^, a hallmark MDP gene specifically demethylated in the myeloid lineage across the hematopoietic system.

To bridge our mechanistic findings with physiological relevance, we systematically validated our results in primary hematopoietic and AML datasets. Despite distinct cellular origins, we observed striking DNAm similarities at TET2 binding sites between *in vitro* and *in vivo* myeloid commitment. This analysis uncovered a previously unreported intragenic enhancer within the *AGO2* locus, where TET2-driven demethylation and *AGO2* activation dynamics closely tracked hematopoietic maturation toward the myeloid lineage. While previous studies have implicated AGO2 in granulocytic, monocytic, and erythroid differentiation ^31–34,41^, our functional data reveal that AGO2 depletion limits proliferation, promotes myeloid differentiation, and abrogates leukemic engraftment *in vivo*. These findings position AGO2 as a central regulator of hematopoietic cell fate and leukemia progression. However, the molecular events underlying these effects warrant further investigation, particularly given AGO2’s multifaceted regulatory potential—encompassing canonical RISC-dependent and non-canonical functions ^31,32,42–44^, as well as emerging roles in mRNA sequestration within P-bodies ^45,46^, a process recently linked to myeloid leukemia ^47^.

Overall, our study uncovers a TET2-dependent regulatory mechanism at the *AGO2* locus, whereby lineage-specific enhancer demethylation fine-tunes *AGO2* expression to support proper myeloid differentiation, while its disruption may lead to leukemogenesis. Beyond mechanistic insight, these findings highlight AGO2 as a potential therapeutic target in AML, either through direct pharmacologic inhibition ^48^ or in combination with DNA hypomethylating agents, offering a strategy to restrain leukemic growth while preserving normal hematopoietic function.

## METHODS

The research conducted in this study complies with all relevant ethical regulations. All experimental procedures were approved by the Animal Care Committee of the Barcelona Biomedical Research Park (DAAM 9667/DAAM11883).

### Mice Care and Maintenance

7- to 16-week-old non-obese diabetic/NOD.Cg-Prkdcscid Il2rgtm1Wjl Tg(CMV-IL3,CSF2,KITLG)1Eav/MloySzJ (NSGS) mice (The Jackson 506 Laboratory, Ban Harbor, ME; RRID: IMSR_JAX:013062), housed under pathogen-free conditions, were used as recipients for transplantation experiments. The animal rooms were fully air-conditioned with a temperature of 20 - 24 °C and 45 – 65% humidity, according to Annex A of the European Convention EC. The maximum stocking density of the cages corresponds to Annex III of the 2010/63 EU. The cages were constantly filled with structural enrichment and the animals had unlimited access to food and water.

### Cell lines growth and maintenance

BLaER1 cells (BlaER) were derived from a human B-cell precursor leukemia cell line (RCH-ACV) that stably expresses the myeloid transcription factor C/EBPα fused with the estrogen receptor (ER) and labeled with GFP ^21^. BlaER cells were grown in suspension in RPMI 1640-HEPES (GIBCO) supplemented with 10% heat-inactivated FBS (GIBCO), 1% Penicillin-Streptomycin (GIBCO), 1% L-glutamine (GIBCO) and 550 µM β-mercaptoethanol (GIBCO). Culture medium was replaced every 2 - 3 days maintaining the cellular density between 0.2 - 2 × 10^6^ cells per ml.

HEK-293T cells were grown in Dulbecco’s Modified Eagle’s Medium (DMEM) (+) D-glucose supplemented with 10% heat-inactivated FBS (GIBCO), 1X Penicillin-Streptomycin (GIBCO) and 1X L-glutamine (GIBCO).

AML-579 PDX cells ^38,39^ were cultured in RPMI 1640 (GIBCO) supplemented with 20% heat-inactivated FBS (GIBCO), 5% L-glutamine (GIBCO), 1% Gentamycin (GIBCO), 1% Penicillin-Streptomycin (GIBCO), 0.6% Insulin-Transferrin-Selenium (Life Technologies, cat. no. 41400045), 1 mM Sodium Pyruvate (GIBCO) and 550 µM β-mercaptoethanol (GIBCO).

For optimal growth, all cell lines were kept in a 5% CO2 humidified atmosphere at 37 °C. Cells were checked for mycoplasma infection every month and tested negative.

### In vitro myeloid commitment of human B cells into induced macrophages

To induce myeloid commitment of leukemic BlaER B cells (B-cell) into induced macrophages (iMac), cells were seeded at 0.2 × 10^6^ cells per ml in medium supplemented with 100 nM β-estradiol (E2) (Sigma Aldrich) to activate C/EBPα. Additionally, human IL-3 (10 ng ml^−1^; PeproTech) and M-CSF (10 ng ml^−1^; PeproTech) were added to favor the conversion.

### Assessment of in vitro myeloid commitment by flow cytometry

0.1 × 10^6^ cells were blocked with human Fc Receptor binding inhibitor (eBiosciences) for 10 min, followed by the addition of conjugated CD11b (CD11b-APC BD #550019) and CD19 (CD19-PE BD #340364). Antibody staining was performed in the dark for 20 min at room temperature. Prior to the acquisition, cells were washed once with PBS and stained with DAPI (4’,6-diamidino-2-phenylindole) for viability assessment. Sample acquisition was performed using the BD FACSCanto II Flow Cytometry System (BD Biosciences) and data analysis with the FlowJo software.

### TET2 gene editing vectors construction

The TET2-3×FLAG-P2A-NeoR/HygR targeting vectors were cloned by serial modification of base vectors pMK292 (Addgene #72830), pMK293 (Addgene #72831) and pFETCh Donor (Addgene #63934). Briefly, 800 bp homology arms (5’ HA and 3’ HA) at the last coding exon of *TET2* were amplified from BlaER genomic DNA. pMK backbone, NeoR/HygR and 3×FLAG-P2A elements, were amplified from pMK293, pMK292 and pFETCh donor plasmids respectively. All fragments were assembled by Gibson reaction ^50^ for 1 hour at 50 °C. Gibson assembly master mixes were prepared by the Protein Technologies Unit at the Centre for Genomic Regulation (CRG, Barcelona). Guide RNAs (gRNAs) targeting TET2 last coding exon were cloned into px330-mcherry (Addgene #98750) empty backbone following Zhang lab guidelines (https://www.zlab.bio/resources).

### shRNA knockdown vectors construction

For doxycycline inducible shRNA-mediated TET2 silencing, oligonucleotide pairs encoding shRNAs were annealed and cloned into Tet-pLKO-puro plasmid (Addgene, # 21915). The following shRNAs were used:

shCTRL: GTGGACTCTTGAAAGTACTAT

shTET2.1: TTTCACGCCAAGTCGTTATTT

shTET2.2: CAGTCTAATGTACGAACTTTA

shTET2.3: CAGATGCACAGGCCAATTAAG

For constitutive shRNA-mediated *AGO2* silencing, oligonucleotide pairs encoding shRNAs were annealed and cloned into pSICOR-PGK-puro (Addgene, #12084) or modified pSICOR-BFP-puro (from Addgene, #31845) plasmids. The following shRNAs were used:

shCTRL: GTGGACTCTTGAAAGTACTAT

shAGO2.1: GCAGGACAAAGATGTATTA

shAGO2.2: GCAGGACAAAGATGTATTA

### Generation of TET2-FLAG epitope tag knock-in

To generate TET2-3×FLAG knocked-in cells, two rounds of transfection were performed with the TET2-3×FLAG-P2A-NeoR and TET2-3×FLAG-P2A-HygR plasmids to target both TET2 alleles. Transfection was carried out by electroporation with Amaxa Nucleofector KitC (Lonza) where 1 µg of TET2-3×FLAG-P2A-NeoR plasmid per 100 µl of Nucleofector solution mix, was used to electroporate 1 × 10^6^ BlaER cells (Nucleofector II Device, Program: X-001). Two days after transfection, Neomycin at 500 µg ml^−1^ (Sigma-Aldrich) was added to the medium, which was changed every 2–3 days until cells recovered normal proliferation rates. Neomycin-resistant cells were then subjected to the same transfection procedures using the TET2-3xFlag-P2A-HygR plasmid to generate double-resistant cell lines selected with hygromycin at 100 µg ml^−1^ (Gibco). The bulk population resistant to the drug was sorted by FACS to generate single-cell clones. After 2 weeks of expansion, homozygous insertion was confirmed by genotyping PCR with the following primers:

Forward: CATGAAACTTCAGAGCCCACTTAC

Reverse: AGATAACCTCTTTTGTTGCTGGTG

### Cellular transduction for shRNA-mediated silencing

shTET2-Tet-pLKO-puro and shAGO2-pSicoR-puro BlaER cell lines and their respective controls were generated by lentiviral transduction. For each plasmid, low-passaged HEK293T cells were seeded at 0.3–0.5 × 10^6^ cells per ml and co-transfected with VSV-G and psPAX2 lentiviral plasmids using a calcium phosphate-mediated transfection protocol. Medium change was performed 12 hours post-transfection and supernatants containing lentiviral particles were harvested and filtered (0.45 μm) at 48 hours post-transfection. Viral particles were concentrated at 70,000 × g for 2 hours at 10 °C, and then the viral pellets were resuspended by agitation for 3 hours at 4 °C. Then, 1 × 10^6^ BlaER cells were spin-infected (1,000 × g) at 32 °C for 90 minutes and 12 hours post-infection cells were washed to remove debris and remaining viral particles.

Two days post-infection, puromycin at 1 µg ml^−1^ was added to the medium and maintained for 3-7 days to select the transduced cells. Selection medium was changed every 2–3 days.

For lentiviral transduction of AML-579 PDX cells, shAGO2-pSicoR-BFP or shCTRL-pSicoR-BFP viral supernatants were collected 24 and 48 hours after transfection and concentrated by centrifugation at 70.000 × g for 2 hours at 10 °C. Concentrated virus was resuspended in PBS and stored at -80 °C. Cells were transduced by spin-infection with virus at 1,000 × g for 90 min at 32 °C, in the presence of Polybrene (8 µg ml^−1^).

### Xenotransplantation of primary human AML

For leukemic engraftment evaluation in AGO2-depleted conditions, AML-579 PDX-eFFly-mCherry cells were transduced with shAGO2-pSicoR-BFP and shCTRL-pSicoR-BFP (see: Cellular transduction for shRNA-mediated silencing). 48 hours post transduction, cells were T cell-depleted with an anti-CD3-OKT3 antibody (LABCLINICS) and FACS sorted to reach a purity of more than 95% of BFP^+^ cells. For transplantation, sub-lethally irradiated (2 Gy) NSGS mice were subjected to intra-tibial injection with 0.1 × 10^6^ PDX cells (n = 4 per group).

### AML PDX graft evaluation

shAGO2 and shCTRL PDX tumor burden was monitored bi-weekly by bioluminescence (BLI) (starting from 2 weeks post-injection) using Xenogen *in vivo* imaging system (IVIS) (Perkin Elmer). Data was quantified with Aura Imaging Software.

Mice were euthanized when disease symptoms were evident, leukemia engraftment was incompatible with animal welfare (Max. Radiance > 108 photons/s/cm2/sr), body weight reduction of 20% or at experimental endpoint (26 weeks post-injection).

At 10 weeks post-injection, when the first shCTRL mouse was euthanized, peripheral blood (PB) was collected by facial vein bleeding and bone marrow (BM) by aspiration from non-injected tibiae. The leukemic engraftment was analyzed by flow cytometry. To remove erythrocytes, PB and BM aspirates were lysed with BD FACS TM Lysing solution (BD Biosciences cat. no. 349202) for 15 min. Next, to detect surface markers, cells were first incubated with anti-mouse CD16/CD32 Fc Block (BD Biosciences cat. no. 553141) for 10 min on ice to prevent background staining. Cells were then stained with APC-anti-human-HLA-ABC (BD Biosciences cat. no. 555555) antibody on ice, protected from light, for 20 min. Next, cells were washed and acquired using a BD LSRFortessa SORP (BD Biosciences). Human blasts were defined as HLA^+^/mCherry^+^ cells.

At the endpoint, all mice were analyzed for leukemic grafts in BM, liver, and spleen tissues, which were isolated after euthanasia and analyzed by flow cytometry as previously described.

### TET2 knockdown induction with doxycycline

To induce TET2 KD in shTET2-Tet-pLKO cell lines, shRNA activity was triggered by the addition of doxycycline (Dox) at 1 µg ml^−1^ (Sigma) to the medium. Dox was added 48 hours prior to myeloid commitment induction and re-added every 2 days during the process to maintain a sufficient concentration for shRNA expression at all timepoints. TET2 reduction was confirmed by RT-qPCR.

### Western blots

For protein extraction, up to 10 × 10^6^ cells were incubated in lysis buffer (50mM ph7.5 Tris-HCl, 150mM NaCl, 1mM EDTA, 1mM EGTA, 5mM Mg2Cl, 0.5% Triton X-100 and proteinase inhibitors) on ice for 1 hour, followed by centrifugation at 12,000 × g for 15 min to remove cell debris. Up to 100 μg of clean protein extract was separated by electrophoresis in an 8% polyacrylamide gel and transferred to a nitrocellulose blotting membrane at 400mA for 4 hours. Membranes were blocked with 5% nonfat milk in TBS– Tween (50 mM Tris, 150 mM NaCl, 0.1% Tween 20) for 30 min, shaking at room temperature. Primary antibodies incubation was done at 4 °C overnight, followed by secondary antibodies staining for 1 hour at room temperature and protected from light. The membranes were visualized using Odyssey CLx Imager (LI-COR).

The following antibodies were used: rabbit anti-human TET2 (1:1,1000 dilution) (Abcam, cat. no. ab94580), rabbit anti-human AGO2 (1:1,000 dilution) (Cell Signaling, cat. no. C34C6), mouse anti-human α/β-Tubulin (1:5,000 dilution) (Cell Signaling, cat. no. 2148), goat anti-rabbit Alexa Fluor 680 (1:10,000 dilution) (Invitrogen, cat. no. A21109), goat anti-rabbit Alexa Fluor 790 (1:10,000 dilution) (Invitrogen, cat. no. A11375).

### Intracellular Flow Cytometry Staining

For 5hmC and AGO2 intracellular staining, cells were fixed with 4% methanol-free formaldehyde for 15 min at room temperature following permeabilization and blocking with 0.1% Triton-X containing 2% donkey serum. For 5hmC staining, DNA was denatured with 2N HCl for 30 min and neutralized with Tris-HCl (pH 8.0) prior to the blocking step. Cells were incubated with the primary antibodies at room temperature for 2 hours for 5hmC or at 4 °C for 1 hour for AGO2, followed by fluorochrome-conjugated secondary staining for 30 min and protected from light.

The following antibodies were used: rabbit anti-human 5hmC (1:500 dilution) (Active Motif, cat. no. 39769), mouse anti-human AGO2 (1:400 dilution) (Proteintech, cat. no. 67934-1-Ig), goat anti-mouse Alexa Fluor 647 (1:400 dilution) (Invitrogen, cat. no. A21235), goat anti-rabbit Alexa Fluor 647 (1:400 dilution) (Invitrogen, cat. no. A21245).

### Proliferation assays

shAGO2 and shCTRL cells were plated at 0.2 × 10^6^ cells and monitored by flow cytometry for proliferation in either uninduced or myeloid conversion conditions with CellTrace Violet (Invitrogen Cat #C34570). 7-Aminoactinomycin D (7-AAD) was used as a viability dye).

Manual counting was done in uninduced B-cells at 5 days post-seeding with a hemocytometer (Trypan Blue Exclusion).

### Quantitative RT-PCR

Total RNA was extracted using Trizol (eBioscences) and 0.5 µg of total RNA was converted into cDNA using the High-Capacity cDNA Reverse Transcription Kit (Applied Biosystems). Real-time quantitative PCR reactions were performed using SYBR Green Universal Master Mix (Applied Biosystems) and analyzed using QuantStudio 5 System (Applied Biosystems). *HPRT1* was used as an internal control. The following primer pairs were used:

HPRT-F: GACCAGTCAACAGGGGACAT

HPRT-R: CTGCATTGTTTTGCCAGTGT

TET2 -F: TACCGAGACGCTGAGGAAAT

TET2-R: ACATGCTCCATGAACAACCA

AGO2-F: CAAGTCGGACAGGAGCAGAAAC

AGO2-R: GACCTAGCAGTCGCTCTGATCA

### TET2-FLAG Chromatin immunoprecipitation (ChIP) – seq

TET2 binding sites were evaluated by FLAG-based immunoprecipitation in TET2-3×FLAG-tagged versus untagged control cells. 50 × 10^6^ cells were collected at 72 hours post-induction with 2 biological replicates per condition and washed twice in ice-cold PBS. Following a dual cross-linking ChIP protocol (modified from Broome et al. ^51^), a fresh stock solution of 0.25 M disuccinimidyl glutarate (DSG) (Fisher Scientific) in DMSO was prepared and diluted to 2 mM in PBS. Cell pellets were resuspended in 5 ml of DSG crosslinking solution and incubated in rotation at room temperature for 30 min followed by two PBS washes. Next, the samples were dual crosslinked with 1% formaldehyde (Sigma) in rotation for additional 10 minutes at room temperature. To quench the reaction, 0.125M glycine was added and samples were incubated in rotation for 5 min at room temperature.

Crosslinked cells were washed twice in ice-cold PBS and resuspended in cold IP buffer (1 volume SDS buffer (100mM NaCl, 50mM pH8.1 Tris-HCl, 5mM pH8 EDTA, 0.2% NaN3, 0.5% SDS) : 0.5 volume Triton dilution buffer (100mM NaCl, 100mM pH8.6 Tris-HCl, 5mM pH8 EDTA, 0.2% NaN3, 5% Triton X-100) supplemented with proteinase inhibitors (Roche). Chromatin was sheared to 100 - 300 bp fragments using a Bioruptor Pico Sonicator (Diagenode) at 4 °C for 13 cycles (30s on / 30s off) in 15 ml Bioruptor Pico Tubes (Diagenode). The sonicated chromatin was spun down at 20,000 × g for 20 min at 4 °C to remove any insoluble portion, and the supernatant was pre-cleared with 20 μl of Dynabead A/G mix (Invitrogen). All chromatin was incubated with 5 μg of Monoclonal anti-FLAG M2 antibody (Sigma) overnight at 4 °C in rotation. The next day, 50 μl of Dynabead A/G mix (1:1) were blocked with BSA (5 mg ml^−1^) and added to the sample for immunoprecipitation for 4 hours at 4 °C in rotation.

Chromatin-antibody-bead complexes were washed three times with ice-cold low salt buffer (50mM pH7.5 HEPES, 140mM NaCl, 1% Triton X-100) and once with ice-cold high salt buffer (50mM pH7.5 HEPES, 500mM NaCl, 1% Triton X-100). After the last wash, the complexes were de-crosslinked in elution buffer (1% SDS, 0.1M NaHCO3) by overnight incubation at 65 °C with shaking at 1300 rpm. The next day, the eluted portion was treated with RNase A (20 ng ml^−1^) for 1 hour at 37°C followed by proteinase K (200 ng ml^−1^) for 2 hours at 65 °C. The immunoprecipitated DNA was purified by phenol:chloroform:isoamyl alcohol (25:24:1) extraction.

For ChIP-seq, samples were subjected to quality control and quantification with Agilent 2100. Library preparation and sequencing was performed by an external sequencing service provider using a DNABSEQ-G400 (MGI Tech) sequencer.

### RNA-seq

2 × 10^6^ shTET2 and shAGO2 with their respective shCTRL cells were collected at 0 (B-cell), 72 and 144 (iMac) hours post myeloid induction, with at least 2 biological replicates per group. RNA was extracted with the RNeasy Mini Kit (QIAGEN Cat#74104) and subjected to quality control with Agilent 2100 to ensure high RNA integrity (RIN > 7). For library preparation, rRNA removal was done either with poly-A enrichment (shTET2 RNA-seq) or the RNase H (shAGO2 RNA-seq) following the DNABSEQ Eukaryotic Strand-specific mRNA pipeline. Library preparation and sequencing were performed by the external sequencing service provider using DNABSEQ-G400 (MGI Tech) sequencer.

### DNA methylation arrays

1 × 10^6^ shTET2 and shCTRL cells were collected at 0 (B-cell), 72 and 144 (iMac) hours post myeloid induction, with 3 biological replicates per group. DNA was extracted with Wizard Genomic DNA Purification Kit (Promega) and 1 µg of genomic DNA was bisulfite (BS)-converted using the EZ DNA Methylation Gold Kit (Zymo Research). 100-500 ng of bisulfite converted DNA was hybridized to Infinium MethylationEPIC v1.0 Bead-Chip arrays. Hybridization procedures were performed by the Genomics Unit of the Josep Carreras Leukaemia Research Institute following the manufacturer’s instructions.

### DNA methylation validation by pyrosequencing

Bisulfite (BS)-converted DNA from shTET2 and shCTRL cells (see: DNA methylation arrays) was additionally used for pyrosequencing validation of DNA methylation levels at the *AGO2* regions of interest (ER1 and ER2). Converted DNA was PCR-amplified using the IMMOLASE DNA polymerase Kit (Bioline) which product was sequenced with the Pyromark Q48 system (QIAGEN). Primers design and DNA methylation levels analysis was done with PyroMark Assay Design 2.0 software (QIAGEN).

The following amplification primer were used:

ER1-F: TGTGTTAGGTATAGTGTTAGGGGTTTA

ER1-R: AATATAATAACCCCTATACCCCCACAACAT

ER2-F: TGTTTGTTGGAAATTAGGTATATGAG

ER2-R: AACCTTAAAAAAAAAAACCCTATTACTAT

The following sequencing primer were used:

ER1-seq-1: GTAAGAGAAATAAGAAAATAGAAG

ER1-seq-2: AGTAAGGGTTTGTTTGGG

ER1-seq-3: GGGTTAGGGGTTGTG

ER1-seq-4: AGGTATAGTGTTAGGGGTTTAA

ER2-seq-1: ACTAAATCCAAACTCTACC

ER2-seq-2: AAAAACCAACCCAAACAATAAT

### Bioinformatic analyses

All new sequencing and array data obtained were mapped onto the human genome assembly hg38 (Ensembl GRCh38). All bioinformatic analyses were done with Python (v3.11.9) and R environments (4.2.1) using packages from the Bioconductor suite (v3.0) ^52^. Genomic heatmaps, average plots, and bigwig tracks were generated using deepTools (v3.3.1) ^53^. The integration of different datasets was performed using the bedtools package (version 2.31.1) ^54^. Gene Ontology (GO) enrichment analyses were performed using the clusterProfiler (v4.4.4) ^55^. Custom plots integrating GO enrichment with expression data were done with plotGODESeq package (https://gitlab.com/ngamblen/plotGODESeq). Gene Set Enrichment Analysis (GSEA) were done using the fgsea package (v1.32.2) ^56^. Ternary Diagrams were generated with the ggtern package (3.5.0). The MA plots were generated using ggmaplot function from ggpubr R package (v0.6.0). Heatmaps and clustering analyses were performed using the ComplexHeatmap (v2.10.0) ^57^ and deepTools computeMatrix (v3.3.1) tool ^53^. The Principal Component Analysis (PCA) were performed using limma (v3.52.2) ^58^ and prcomp function from the stats package (v4.4.0). Upset plots were generated using the upsetR package (v1.4.0) ^59^. Correlations were calculated with the sm_statCorr from the smplot2 R package (v0.2.5). Kaplan–Meier survival analysis plots were done with the survit function from the survival package (v3.8-3) and plotted with the survminer package (v0.5.0). The remaining plots were generated using the R package ggplot2 (v3.4.2).

### TET2 ChIP-seq analysis

Reads were trimmed using the TrimGalore package (v0.6.6) to remove adaptors and mapped using Bowtie2 (v2.4.4.1) ^60^. Duplicated reads were removed with the tool MarkDuplicates from the package Picard (v3.1.0) (Picard Tools https://broadinstitute.github.io/picard). Peaks were called against the control samples using MACS2 (v2.2.5) ^61^, parameter (-q 0.05). For peak calling, regions overlapping the ‘Encode blacklist’ regions were removed ^62^, as well as mitochondrial reads. Peaks were annotated to genomic features in R with the package ChIPseeker (v1.32.1) ^63^. Biological replicates were handled with Irreproducibility Discovery Rate (IDR) method ^64^, using the package idr (v1.3).

### Hi-C analysis

Raw Hi-C interactions at 0 (B-cell), 72 and 144 (iMac) hours post induction were processed with the HiC-Pro (v3.1.0) ^65^ pipeline following the recommended parameters with ICE normalization. Normalized contact maps at 5 Kb resolution were used for downstream analysis.

The matrices were normalized and corrected with the hicNormalize function to the lower sequencing depth (--smallest) and hicCorrectMatrix function, respectively, from the HiCExplorer package (v3.7.6) ^66^. Publicly available Hi-C datasets from healthy hematopoiesis were converted from HiC ‘.pairs’ format to 5 Kb binned cool files with the cooler package (v0.10.3) and normalized as previously described. Contacts heatmap was plotted with the HiContacts package (v3.21) ^67^.

### Identification of TET2 bound regulatory regions

For Cis Regulatory Elements (CREs) definition, intersecting of ATAC-seq peaks with H3K4me1 and transcription start sites (TSS) was done as previously described ^23^. The evaluation of H3K27ac signal was done with the Diffbind package using the DBA_DESEQ2 method (v.2.2.12). H3K27ac loss or gain was defined by comparing iMac vs B-cell timepoints (signal (FC)>2; FDR<0.05). H3K27ac transient gain was defined by an increase signal at 72 hpi versus B-cell and iMac stages (signal (FC)>1; FDR<0.05). TET2 binding sites were overlapped with the defined CREs using the foverlaps function from the data.table package (v1.17.6)

### TET2 enhancer–promoter loops definition

Hi-C data integration with TET2 sites CREs classification (see: Identification of TET2 bound gene regulatory regions) was used to define Enhancer (E) – Promoter (P) sites. Contact intensity between bin pairs where TET2 overlapping occurred on both sites was extracted from ‘bedpe’ files and classified according to their CRE definition (E-P, E-E, P-P). Only the bin pairs where E-P loop occurred were used for the downstream analysis. Gene expression integration considered only the promoter-associated gene for the quantification.

### Long-range interactions

Aggregate Peak Analysis (APAs) showing long-range interactions between TET2 binding sites were computed using the HiCExplorer package (v3.7.6) tool hicAggregateContacts. Hi-C matrices were generated at 5 Kb resolution using HiCExplorer and long-range interactions (5–10 Mb) between the provided coordinates. APAs quantification was done following ENCODE guidelines for Hi-C processing and analysis (https://www.encodeproject.org/documents/75926e4b-77aa-4959-8ca7-87efcba39d79/@@download/attachment/comp_doc_7july2018_final.pdf).

### Whole Genome Bisulfite Sequencing (WGBS) analysis and quantification

WBGS data during myeloid commitment was processed as previously described ^24^. DNA methylation quantification within TET2 binding sites was calculated from normalized bigWig files with Python numpy.nanmean function (v2.1). Only confident CpGs (>3x coverage) were considered, and changes in DNA methylation equal to or greater than 15% were classified as dynamic.

Methylome data from 30 AML samples with distinct mutational profiles were obtained from previous studies (n=18 ^68^ and n=12 ^69^). Pre-processed bigWig files were used for the downstream analysis. For DNA methylation quantification at the AGO2 locus, only IDR < 0.05 TET2 binding coordinates were considered as regions of interest (R1-7). Coordinates lift-over from hg19 to hg38 (for Spencer et al ^69^) was done with the UCSC Lift Genome Annotations tool (https://genome.ucsc.edu/cgi-bin/hgLiftOver).

Sample IDs are listed in **Table S2**.

### RNA-seq analysis

For shTET2 and shAGO2 RNA-seq datasets, reads were mapped using STAR with standard options (v2.7.6) ^70^. Gene expression was quantified using the function featureCounts from the package Subread (v2.0.3) ^71^. Differentially expressed genes (FDR < 0.05) were detected using the R package DESeq2 (v1.36.0) ^72^. DESeq2 or vsd count normalization was used for further analysis unless stated otherwise Normalized expression for healthy primary hematopoiesis from the Blueprint consortium (http://blueprint-data.bsc.es/#!/about) was obtained from pre-processed RNA-seq data using the RSEM gene-level estimated counts using the R package DESeq2 (v1.36.0) following the tximport() pipeline.

To obtain differentially expressed genes from AML patients, raw counts from the BeatAML2 (https://biodev.github.io/BeatAML2/) and TCGA-LAML cohorts were processed as previously described. BeatAML2 patients’ overall survival, leukemia classification and mutational status were extracted from the Clinical Summary report (https://biodev.github.io/BeatAML2/).

### Transcription factor motif enrichment analysis

TET2 sites motif enrichment analysis was performed using MEME Suite - motif-based sequence analysis tool. Simple Enrichment Analysis (v5.5.7) ^73^ tool was used to determine relative transcription factor enrichment with shuffled input sequences as the background (https://meme-suite.org/meme/tools/sea).

### Transcription factor activity inference

The TFs’ activity estimation (activation or inhibition) was done with decoupleR (v2.14.0) ^74^ using the CollecTRI regulons ^75^. Significant t-values (stat) from differentially expressed genes (iMac vs B-cell) detected in MSPs and MDPs groups were used for the analysis.

### DNA methylation arrays analysis

IDAT raw data from the Infinium MethylationEPIC v1.0 Bead-Chip arrays were processed in R following the minfi pipeline (v1.42.0) ^76^. Probes with low detection p-value (< 0.01), probes with a known SNP at the CpG site, and known cross-reactive probes were removed. For the resulting CpGs, β-values were calculated using minfi functions. Differentially methylated positions (DMPs) were calculated using the limma package (v3.52.2), adjusting by the Benjamini-Hochberg method. Only DMPs with an absolute Δβ-value > 0.15 and p-value < 0.01 were considered for the downstream analysis.

Publicly available Infinium HumanMethylation450 BeadChip IDAT raw data from the TCGA-LAML cohort was processed as previously described. Patients’ overall survival and mutational status was extracted from the cBioPortal database (https://www.cbioportal.org/) ^77,78^. Sample IDs are listed in **Table S3**.

### Statistics and reproducibility

Details for statistical analyses, including the number of biological replicates and/or sample size, are provided in the figure legends. Statistics were calculated using the R stats package (v4.4.0). Data normality was tested before conducting statistical analyses.

In figures, all box plots display the first and third quartiles as the lower and upper edges of the box, with a line inside the box showing the median value and whiskers extending to 1.5 times the interquartile range. An unpaired two-tailed Student’s t-test was used to determine statistical significance in a two-sided manner, unless otherwise specified. Error bars represent standard deviation. When comparing gene expression, ‘differential gene expression’ statistics are shown (see: RNA-seq analysis).

## ACKNOWLEDGMENTS

We thank all members of the Sardina group for their critical reading of the manuscript and for stimulating scientific discussions. We thank T. Graf for kindly sharing the cellular model of myeloid commitment. We thank the IJC-IGTP Flow Cytometry Unit for assistance with setup and cell acquisition, and the IJC Genomics Unit for technical assistance with processing the MethylationEPIC arrays.

We thank CERCA Programme/Generalitat de Catalunya and Fundació Josep Carreras-Obra Social la Caixa for institutional support.

## AUTHOR CONTRIBUTIONS

A.L. designed, performed and analyzed cell culture and molecular biology experiments, designed and performed bioinformatic analyses and prepared figures. G.V., M.O., C.B. and C.F. performed additional cell culture and molecular biology experiments. A.V.L-R and G.S. assisted with bioinformatic analyses. G.V., A.M. and P.M. performed and analyzed the AML patient-derived xenograft experiments. J.J.R.-S. provided essential clinical expertise and contributed to the interpretation and translational contextualization of the results. I.J. provided the luciferase-expressing AML-PDX cells and key experimental data. J.L.S. conceived the study, secured funding, supervised the research, and prepared figures. A.L. and J.L.S. wrote the manuscript with input from all co-authors.

## DATA AVAILABILITY

RNA-seq, ChIP-seq and Infinium MethylationEPIC arrays data generated in this study have been deposited in the Gene Expression Omnibus dataset under accession code GSE307340 (token: xxxxxx). Other *in vitro* myeloid commitment datasets analyzed in this study are available under the following accession numbers: WGBS (GSE271842); ATAC-seq, CEBPA ChIP–seq, TT-seq (GSE131620); RNA-seq, Hi-C, and CTCF ChIP-seq (GSE140528). The H3K27ac, H3K4me1, and H3K4me3 ChIP–seq datasets are available in the ArrayExpress database under the accession number E-MTAB-9825.

The WGBS and RNA-seq datasets across primary human blood cell populations analyzed in this study were generated by the Blueprint Consortium and are available through the Ensembl project (https://projects.ensembl.org/blueprint/) via its FTP site for processed data. Hi-C data on primary HSPC and primary monocytes and macrophages are available under the accession numbers GSE152135 (GEO database) and E-MTAB-10848 (ArrayExpress database), respectively.

The human AML DNAm array data were derived from the TCGA Research Network (http://cancergenome.nih.gov/) and RNA-seq from BeatAML2 repository (https://biodev.github.io/BeatAML2/). The collection of WGBS data in primary AML samples is available under the accession number GSE152099 and at the following URL: https://wustl.app.box.com/s/6fvifntz6golh4vu52epkfqxeyd0vqmk/folder/11145725870. *AGO2* expression during normal hematopoiesis was obtained from the BloodSpot 3.0 database (https://bloodspot.eu/). The “Normal human hematopoiesis (DMAP)” dataset was used to obtain mRNA expression levels of microarray data (log_2_).

## Supplementary Materials

### SUPPLEMENTARY FIGURES

**Figure S1.**
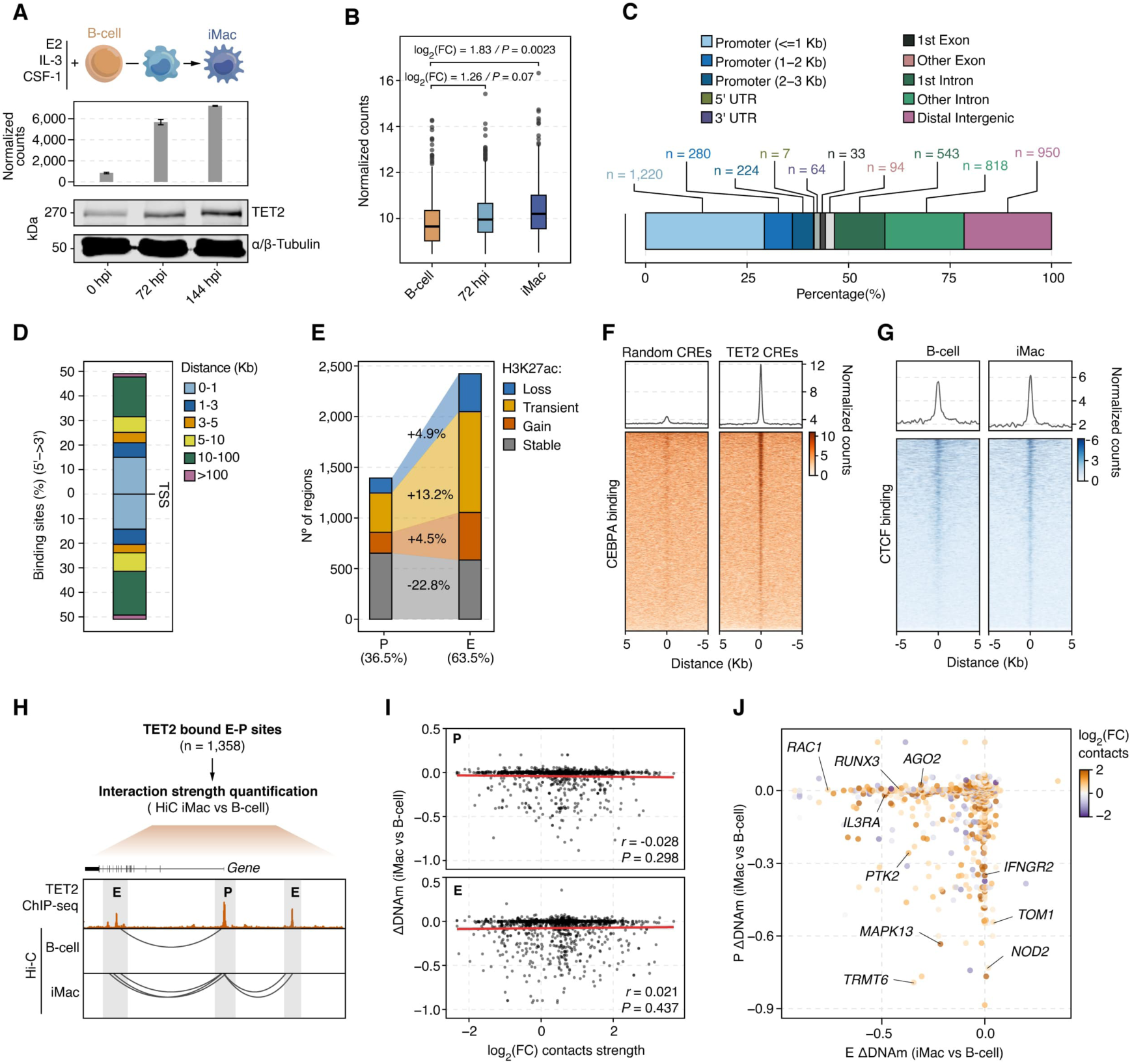
TET2 occupancy links enhancer–promoter architecture to activation of myeloid gene programs. **(A)** Top: Schematic overview of the cellular model used to study myeloid commitment. Middle and bottom: TET2 mRNA and protein levels during myeloid commitment. **(B)** Boxplots displaying levels of highly expressed macrophage genes detected to be significantly changing during myeloid commitment (iMac vs B-cell). Data from BioGPS Macrophage Gene Set (Harmonizome 3.0) (n = 929). **(C)** Genomic features of TET2 binding sites. **(D)** Distribution of TET2 binding sites relative to the nearest Transcription Start Site (TSS). **(E)** Stacked barplots showing chromatin activity dynamics at TET2-bound promoters and enhancers during myeloid commitment, based on H3K27ac changes. Percentage change is normalized to the total number of regions in each group. **(F)** Average plot (Top) and genomic heatmap (Bottom) showing CEBPA occupancy at TET2-bound CREs compared with random CREs in iMac. **(G)** Average plot (Top) and genomic heatmap (Bottom) of CTCF occupancy at TET2-bound CREs in B-cells and iMacs. **(H)** Schematic overview of the strategy used to define TET2-bound enhancer-promoter (E-P) pairs. **(I)** Scatterplots showing the correlation between DNAm and contact strength at promoters (P) and enhancers (E) during myeloid commitment (iMacs vs B-cell). The solid line represents the linear regression fit, and the shaded area denotes the 95% confidence intervals. P values were calculated from a two-sided Pearson correlation. **(J)** Scatterplot depicting changes in DNAm and contact strength at TET2-bound E-P pairs during myeloid commitment (iMac vs B-cell).

**Figure S2.**
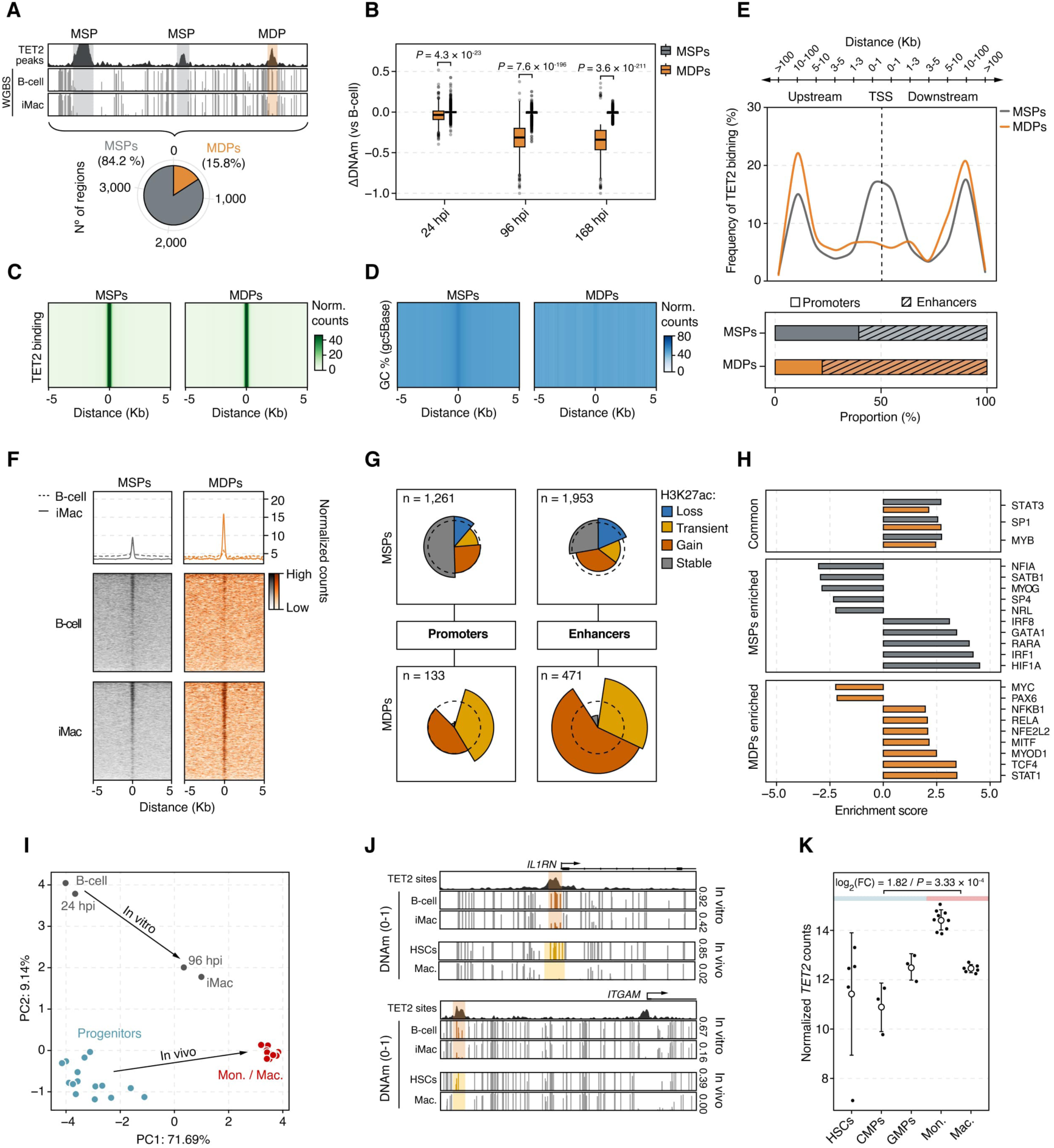
TET2 binding leads to DNA demethylation and activation of a highly dynamic subset of CREs during myeloid commitment. **(A)** Top: Schematic overview of the strategy used to classify TET2-bound sites into Methylation Stable Peaks (MSPs) and Methylation Dynamic Peaks (MDPs). Bottom: Pie chart showing the relative proportions of MSPs and MDPs. **(B)** Boxplots showing DNAm changes (relative to B-cells) during myeloid commitment at MSPs and MDPs. **(C)** Genomic heatmap showing TET2 ChIP-seq signal intensity at MSPs and MDPs. **(D)** Genomic heatmap showing the CG content enrichment at MSPs and MDPs. **(E)** Top: Distribution of MSPs and MDPs relative to the nearest Transcription Start Site (TSS). Bottom: Proportion of the associated CREs at the MSPs and MDPs. **(F)** Average plots (Top) and genomic heatmaps (Bottom) of CEBPA ChIP-seq signal at MSPs and MDPs in B-cells and iMacs. (**G)** Combined pie chart and radar plot showing relative enrichment of TET2-bound CREs and their activation kinetics (H3K27ac levels) within MSPs and MDPs. The radius of each pie segment represents the fold enrichment of MSP and MDP dynamics relative to all CREs, with the dashed line indicating an enrichment value of 1. **(H)** Barplots showing Transcription Factor (TF) activity enrichment inferred from expression changes of genes associated with MSPs and MDPs during myeloid commitment (iMac vs B-cell). Top 10 most enriched (positively or negatively) TF activities are shown for the MSPs group. **(I)** Principal Component Analysis (PCA) of the DNAm dynamics at MDPs during “*in vitro*” myeloid commitment model (B-cell to iMac) versus primary “*in vivo*” (HSCs to macrophages) differentiation. Arrows indicate inferred DNAm trajectories. Progenitors: HSCs: Hematopoietic Stem Cells (n = 9), CMPs: Common Myeloid Progenitors (n = 3), GMPs: Granulocyte-Monocyte progenitors (n = 3). Mon.: Monocytes (n = 9); Mac.: macrophages (n = 6). **(J)** Genome browser snapshot showing example MDPs shared between the *in vitro* and *in vivo* myeloid commitment systems at the *IL1RN* and *ITGAM* loci. Shaded regions highlight TET2 binding sites. Average DNAm levels within each region are shown on the right. **(K)** *TET2* expression (by RNA-seq) dynamics during primary myeloid commitment. Mean ± s.d. are shown. Differential expression analysis was assessed between committed (Mon.;Mac.) and progenitor (HSCs, CMPs, GMPs) cells. Progenitors: HSCs: Hematopoietic Stem Cells (n = 5), CMPs: Common Myeloid Progenitors (n = 3), GMPs: Granulocyte-Monocyte progenitors (n = 3). Mon.: Monocytes (n = 9); Mac.: macrophages (n = 6).

**Figure S3.**
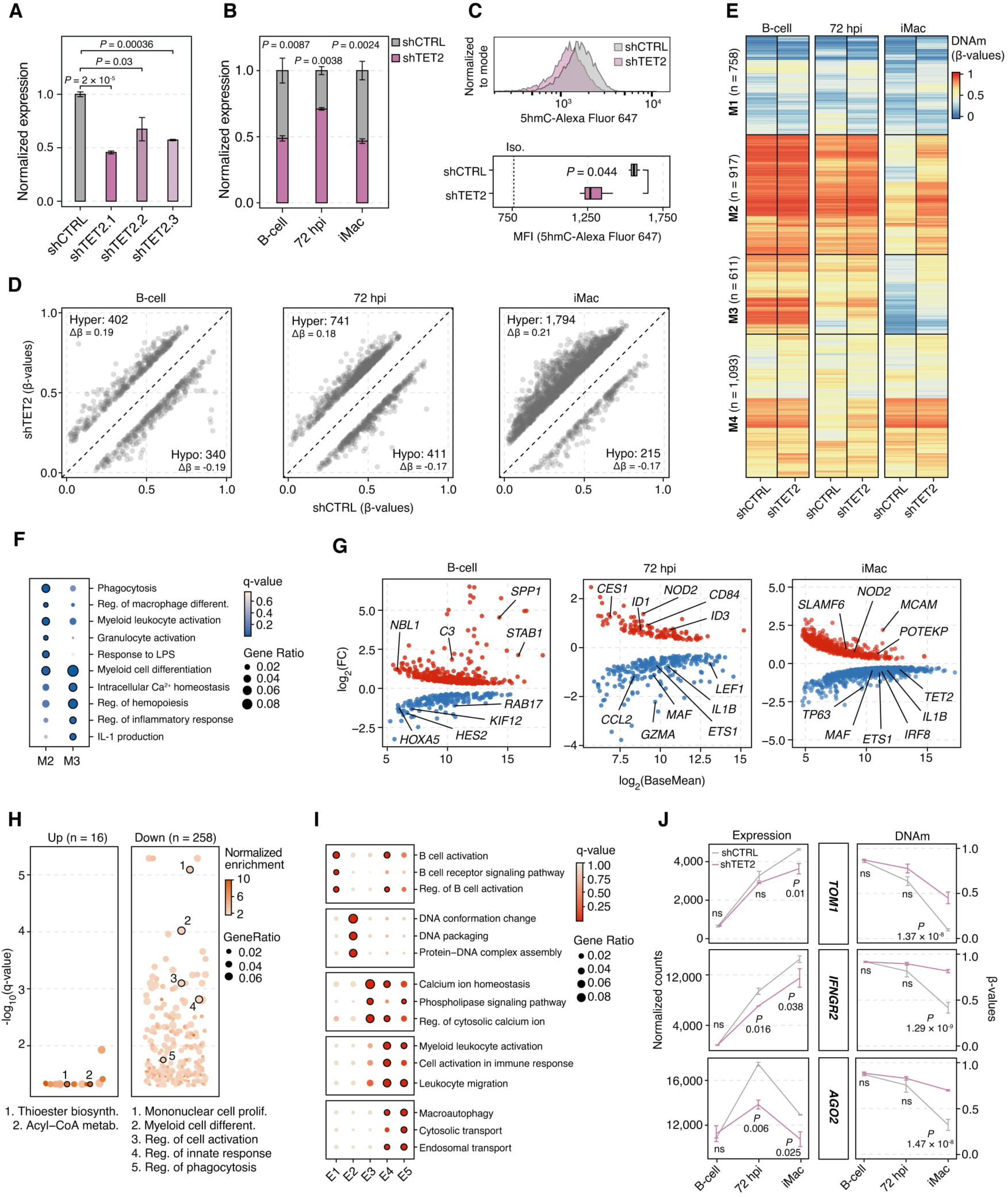
TET2 depletion impairs DNAm-mediated activation of myeloid gene programs. **(A)** Barplots showing TET2 expression (by RT-qPCR) in B-cells transduced with different *TET2*-targeting shRNAs. Values were normalized to the housekeeping gene (*HPRT*) and *TET2* levels in shCTRL cells. Mean ± s.d. are shown for each condition (n = 3). **(B)** Barplots showing TET2 expression (by RT-qPCR) in shTET2 (shTET2.1) and control cells during myeloid commitment. Values were normalized as in **(A)** at each timepoint. Mean ± s.d. are shown (n = 3). **(C)** 5hmC levels (by Intracellular Flow Cytometry) in shTET2 B-cells. Top: representative histograms of 5hmC signal intensity. Bottom: quantification of signal (n = 3 biological replicates). MFI: Mean Fluorescence Intensity. Iso.: Isotype Control. **(D)** Scatterplots showing Differentially Methylated Positions (DMPs) detected in shTET2 versus control cells during myeloid commitment. **(E)** Heatmap showing the clustering of DMPs (M1-M4) detected in shTET2 versus control cells during myeloid commitment. **(F)** Gene Ontology (GO) enrichment analysis of genes associated with M2- and M3-DMP clusters. Selected biological processes from the top 20 most significant terms are shown. Terms with q-value < 0.05 are outlined in black. **(G)** MA plots of Differentially Expressed Genes (DEGs) detected in shTET2 versus control cells during myeloid commitment. **(H)** GO enrichment analysis of the significant up- or downregulated DEGs detected in shTET2 versus control iMacs. Selected biological processes are highlighted with a black outline. **(I)** GO enrichment analysis for the genes associated with expression clusters (E1-E5) from Figure 3F. Selected biological processes from the top 10 most significant terms are shown; significant terms (q-value < 0.05) are outlined in black. **(J)** Expression and DNAm (β-values) levels of representative genes form DEGs/DMPs overlap in shTET2 versus control cells during myeloid commitment. Mean ± s.d. are shown for each timepoint.

**Figure S4.**
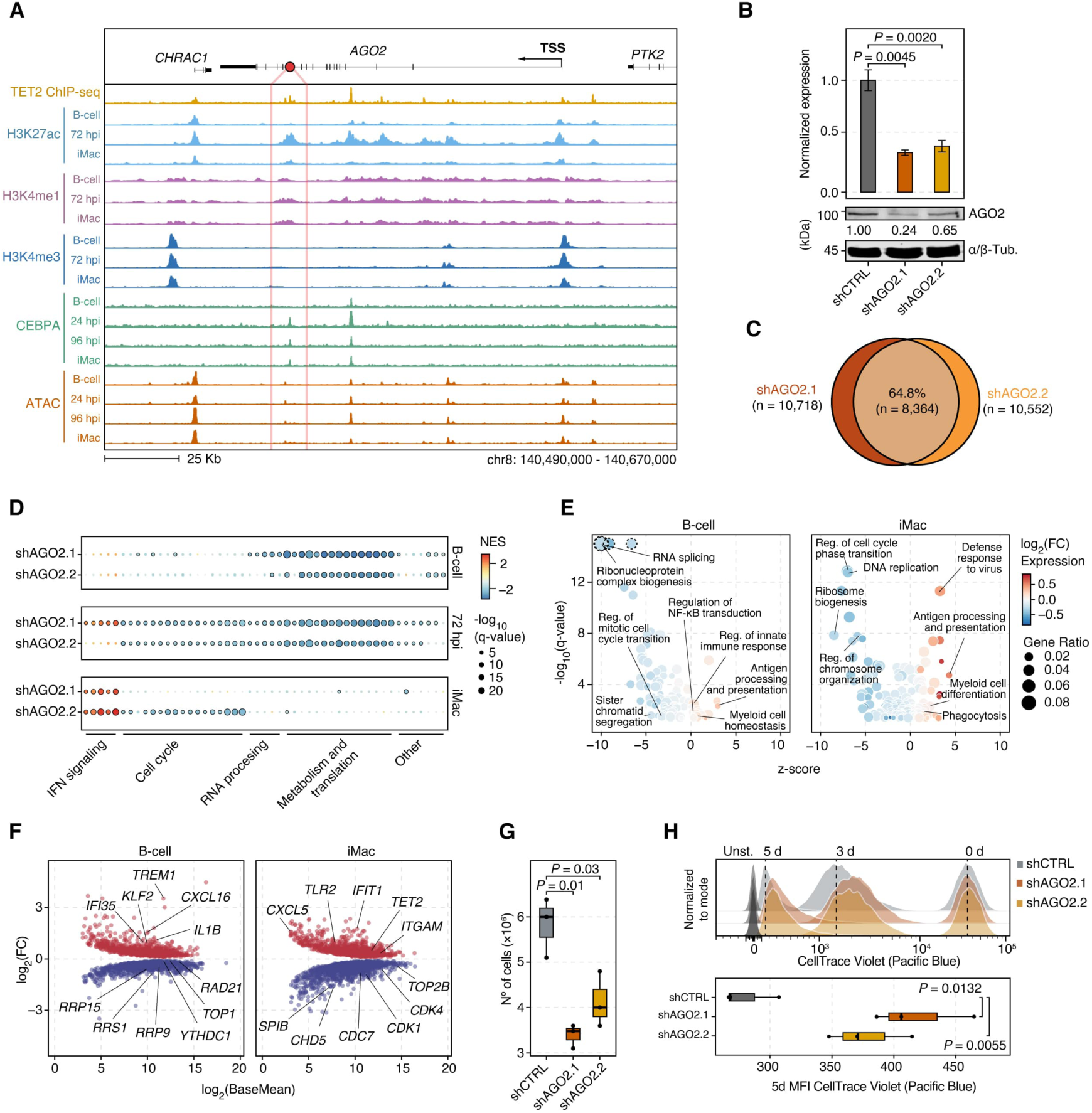
Integrative multi-omics analysis identifies AGO2 as a TET2-dependent regulator of myeloid commitment. **(A)** Genome browser snapshot showing chromatin dynamics and CEBPA-binding at the *AGO2* locus during myeloid commitment. The region of interest identified in Figure 4A is shown as a red dot. **(B)** Evaluation of AGO2 knockdown efficiency. Top: Barplots showing *AGO2* expression (by RT-qPCR) in B-cells transduced with different *AGO2*-targeting shRNAs (shAGO2.1 and shAGO2.2). Values were normalized to the housekeeping gene (*HPRT*) and *AGO2* levels in shCTRL cells. Mean ± s.d. are shown for each condition (n = 3). Bottom: Corresponding AGO2 protein levels normalized to α/β-Tubulin and expressed as a fold change relative to shCTRL. **(C)** Venn-diagram showing the overlap of DEGs detected in the two *AGO2*-targeting shRNAs. **(D)** Gene Set Enrichment Analysis (GSEA) of DEGs in *AGO2*-targeting shRNAs cells during myeloid commitment. Commonly enriched terms from ‘REACTOME’, ‘HALLMARK’ and ‘GOBP’ gene sets are shown and grouped based on their biological function. NES: Normalized Enrichment Score. **(E)** Gene Ontology (GO) enrichment analysis of DEGs in shAGO2 cells at B-cell and iMac stages. Over- and under-represented (z-score) biological processes are plotted, with color indicating the average log_2_(FC) of genes within each term. Outlier terms (- log_10_(q-value) > 15) are indicated by a dotted line. **(F)** MA plots showing DEGs in shAGO2 conditions at B-cell and iMac stages**. (G)** Trypan Blue Exclusion assay showing cell counts of shAGO2 versus control uninduced B-cells at 5 days post-seeding (n = 3 biological replicates). **(H),** Proliferation assay by flow cytometry in shAGO2 versus control uninduced B-cells. Top: representative histograms showing CellTrace Violet signal. Bottom: quantification of CellTrace Violet signal at 5 days post-seeding. Mean ± s.d. are shown for each timepoint (n = 3). MFI: Mean Fluorescence Intensity. Unst: Unstained control.

**Figure S5.**
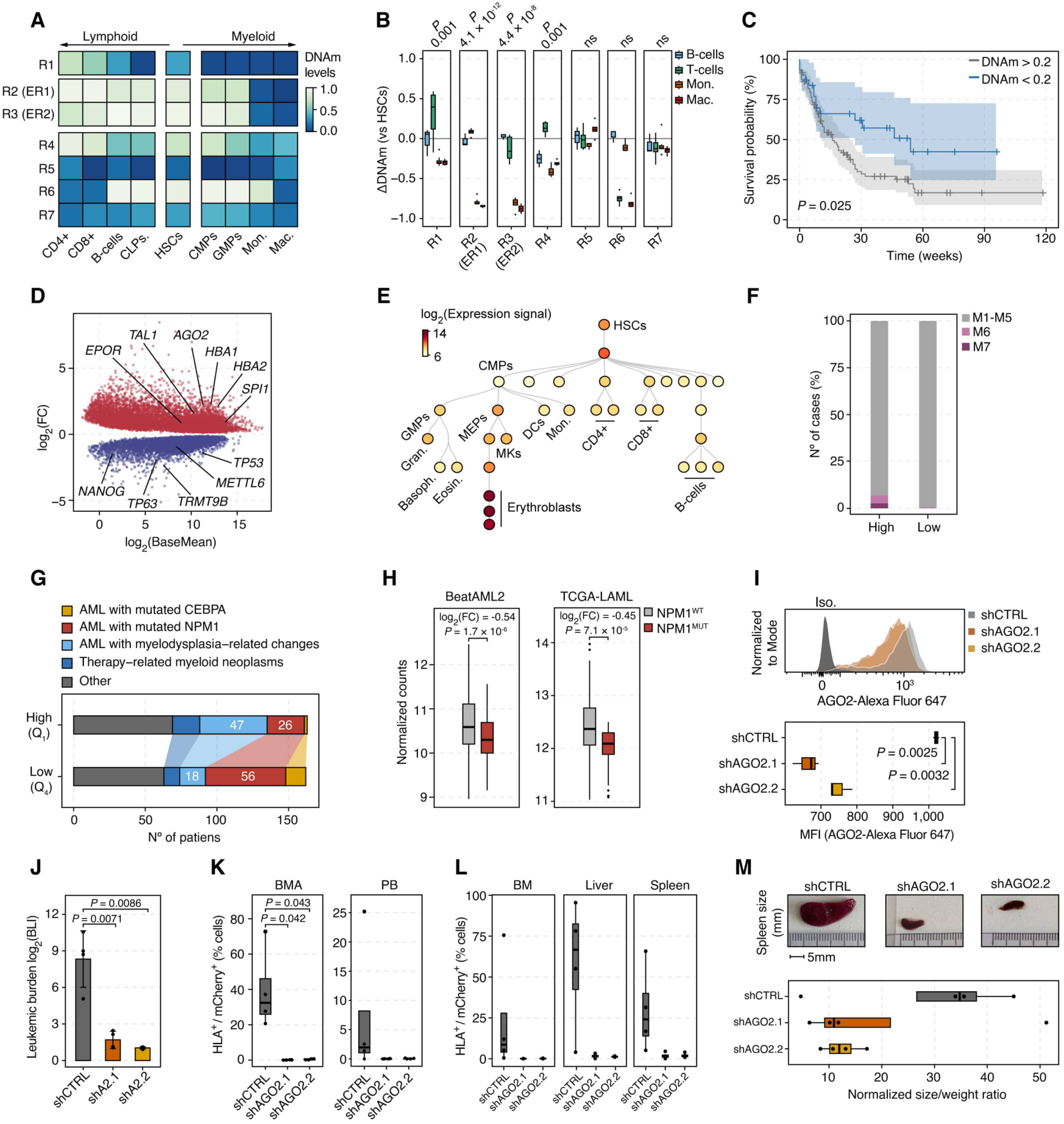
AGO2 is hypermethylated in TET2^MUT^ AML patients and its silencing impairs *in vivo* leukemic progression. **(A)** Heatmap showing DNAm levels at *AGO2* ROIs (R1-R7) during healthy hematopoiesis. **(B)** Boxplots showing the *AGO2* DNAm quantification at ROIs (R1-7) in mature hematopoietic populations. Statistical comparisons were performed between lymphoid (B-cells, CD4+, CD8+) and myeloid (Mon., Mac.) linages. n: HSCs = 9, CLPs = 3, CD8+ = 3, CD4+ = 3, B-cells = 3, CMPs = 3, GMPs = 3, Mon. = 9, Mac. = 6. Mon.; monocytes, Mac.: macrophages. **(C)** Kaplan-Meier survival curves of AML patients from the TCGA-LAML cohort stratified by AGO2 cg00288598 DNAm levels (low: β-value < 0.2, n = 31; and high: β-value ≥ 0.2, n = 95). Shaded areas represent 95% confidence intervals. Statistical significance was calculated using the Mantel-Cox Log-rank test. **(D)** MA plot showing DEGs between patients with high (Q_1_: Quartile 1; n = 163) versus low (Q_4_: Quartile 4; n = 162) *AGO2* expression from the BEAT-AML2 cohort. **(E)** Hematopoiesis tree displaying *AGO2* expression across lineages (data from BloodSpot 3.0) ^79^. **(F)** Stacked barplots showing the proportion of M6 (Acute erythroid leukemia) and M7 (Acute megakaryocytic leukemia) French-American-British (FAB) classified AML subtypes in *AGO2* high (n = 165) versus low (n = 165) expressing patients (BEAT-AML2 cohort). “Other” includes M0-M5, NOS. **(G)** Stacked barplots showing AML subtype distribution at diagnosis in AGO2 high (Q_1_) versus low (Q_4_) expressing patients (BEAT-AML2 cohort). **(H)** Boxplots showing *AGO2* expression in NPM1-mutated versus wild-type AML samples from BEAT- AML2 (left) and TCGA-LAML (right) cohorts. **(I)** AGO2 protein levels (by Intracellular Flow Cytometry) in shAGO2 AML-PDX cells. Top: Representative histograms showing AGO2 signal intensity. Bottom: Quantification (n = 3 biological replicates). MFI: Mean Fluorescence Intensity. Iso.: Isotype Control. **(J)** Barplots of leukemic burden (log_2_(BLI) signal, 10 vs 2 weeks post-injection) in mice engrafted with AML-PDX cells carrying shAGO2 or shCTRL. Mean ± s.d. are shown (n = 4). **(K)** Boxplots showing leukemic engraftment measured by flow cytometry in bone marrow aspirates (BMA) and peripheral blood (PB) from mice carrying shAGO2 or shCTRL AML-PDX cells. **(L)** Boxplots showing leukemic engraftment measured by flow cytometry in organs of mice engrafted with shAGO2 or shCTRL AML-PDX cells at the experimental endpoint. BM: Bone Marrow. **(M)** Spleen size evaluation in mice carrying shAGO2 or shCTRL AML-PDX cells. Top: Representative photos of spleens at euthanasia. Bottom: Boxplots showing the spleen’s weight/size ratio normalized to the animal’s body weight.

